# A modulatory attentional gate promotes recent memory expression in Drosophila

**DOI:** 10.1101/2024.12.20.629828

**Authors:** Ishaan Kapoor, Scott Waddell

**Affiliations:** Centre for Neural Circuits and Behaviour, University of Oxford, Oxford, OX1 3TA, UK

## Abstract

Using short-term memory (STM) requires that animals can distinguish memories of recent experience from those learned previously. In *Drosophila* the neuromodulator octopamine (OA) specifically affects STM through adrenergic-like receptors within neurons in the Mushroom Body (MB) network. However, why OA preferentially impacts STM remains unclear. A fly brain connectome reveals that two OA-VPM3 neurons provide most OA input to the MB network. Artificial OA-VPM3 activation during olfactory learning enhances STM and can divert odor salience. α-adrenergic-like receptor knock-down in MB β and γ lobe MB Output Neurons (MBONs) generally impaired STM, while β-adrenergic-like receptor loss from α lobe MBONs enhanced memory. The identity and connectivity of the relevant MBONs suggests OA modulates an interaction between sites of STM and long-term memory (LTM) storage. Indeed, synthetic, odor, and learning-driven, OA-VPM3 activation temporarily blocks LTM expression. Therefore, OA reconfigures the MBON network so the fly prioritizes expression of recent over remote memories.

## Introduction

Noradrenaline (NA) has been implicated in arousal, alertness and attention in mammals (*1–3*). Broadly projecting noradrenergic neurons from the locus coeruleus (LC) release NA into cortical and subcortical regions of the brain providing an obvious means to coordinate activity across the whole brain, or at least of multiple circuits (*4–6*). NA acts on neurons that express alpha- and/or beta-type adrenergic receptors to affect working memory (*7–10*) and attention (*11–13*). It was therefore proposed that the adrenergic system may direct attention by ‘resetting’ neural networks using ‘Marder-like’ neuromodulation so that they are best configured for the animal’s task in hand (*14*). However, clear examples of such an attentive mechanism have not been identified.

The densely arborizing octopaminergic system of insects is considered a functional equivalent of the mammalian adrenergic system. In *Drosophila,* 27 octopaminergic neuron types project from 4 discrete anatomical clusters to different combinations of brain regions (*15*), which permits an understanding of the functions of identifiable octopaminergic neurons. Importantly, octopamine (OA) modulates target neurons through six G-protein coupled receptors that are sequence homologues of mammalian alpha- and beta-adrenergic receptors (*16*).

OA signaling in the fly has been associated with transient neural processes, such as short-term memory (*17–20*), and several state-switching functions. OA plays a role in fight initiation (*21–22*), sleep-wake transition (*23*), and appetitive persistence-withdrawal (24). OA is also critical for motivational and behavioral state-dependent physiological adaptations (*25–29*). While its role in momentary behaviors suggests that OA, like NA, might represent an attentional signal, the extent of OA’s role in memory and involvement of specific OA neurons remains unresolved.

OA’s roles in short-term olfactory memory and state-dependent odor-tracking involve signaling through neurons in the fly’s Mushroom Bodies (MBs). The MBs are a bilaterally symmetric brain region comprised of around 2,000 (per hemisphere) largely parallel arrays of intrinsic neurons, called Kenyon Cells (KCs), whose sparse activity encodes odor information. Appetitive learning assigns reward value to odors through specific, anatomically-restricted dopaminergic neurons (DANs) which direct plasticity of synapses between odor-activated KCs and the dendrites of particular Mushroom Body Output Neurons (MBONs). OA represents the sweet taste of sugars to reinforce STM via the α-adrenergic-like OAMB receptor in γ4 and β’2a DANs and contributes to the hunger-dependence of learning through the β-adrenergic Octβ2R in PPL101-γ1pedc DANs (*18, 30*). Interestingly, release of OA from feeding-activated OA-VPM4 neurons inhibits odor tracking by limiting the function of MBON γ1pedc, whose dendrites overlap the presynaptic field of PPL101 DANs (*24, 31*). Here we investigated whether OA might modulate memory expression through a network-reset mechanism.

## Results

### OA-VPM3 is broadly presynaptic to neurons of the mushroom body network

The anatomy of MB-innervating OA neurons in the adult fly brain has been described at light microscope resolution of single cells (*15, 18*). We therefore queried their synaptic connectivity within the MB network using the FlyEM hemibrain and FlyWire connectomes (*32–34*) (Figure. 1A). MB-innervating OA neurons primarily arise from two locations in the brain: the ventral midline or the anterior-superior midline. Those with somata in the middle of the ventral side of the suboesophageal zone are either single neurons (Ventral Unpaired Medial, VUM) or bilaterally symmetrical pairs (Ventral Paired Medial, VPM). The Anterior Superior Medial (ASM) neurons have somata in the paired anterior medial clusters among dopaminergic neurons. OA-VPM3 is a single pair of neurons (FlyWire ID: SAD.FB.1, Figure. 1B) that make the greatest number of synapses with KCs, DANs and MBONs (Figure. 1 A-C). OA-VUMa2 predominantly synapses onto KCs in each MB calyx. OA-VPM4, like OA-VPM3, sends several projections to different MB neurons (Figure. S1A). while OA-VUMa6, and OA-VUMa7 mostly project to non-KC targets (Figure. 1A).

**Figure. 1.**
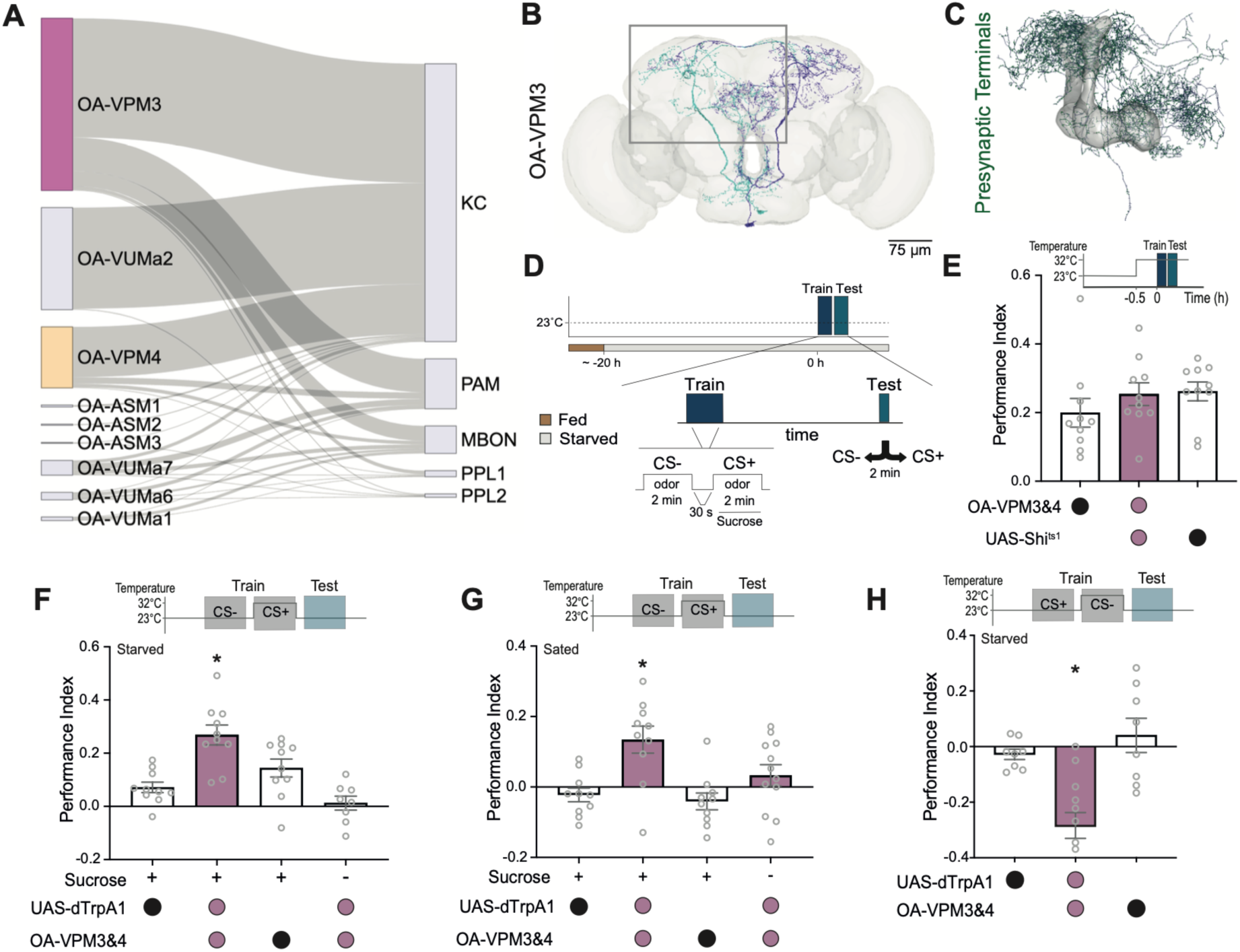
OA-VPM3 activation directs attention to the current odor. (**A**) Sankey diagram from hemibrain connectome of connections between identified OA neurons and numbers of relevant neurons of the MBs. (**B**) Skeleton of OA-VPM3 neuron in the Flywire connectome, box indicates approximate region within the hemibrain connectome. (**C**) OA-VPM3 skeleton from hemibrain connectome with putative presynaptic sites indicated in green. (**D**) Protocol for appetitive olfactory learning and memory testing. (**E**-**H**) Temperature shifting protocols shown above all panels. (**E**) Blocking OA-VPM3&4 (MB022B) output does not affect appetitive STM. **F.** Artificial OA-VPM3&4 activation during odor-sucrose pairing enhances STM. OA-VPM3&4 activation with odor does not substitute for sugar reward. (**G**) OA-VPM3&4 activation during odor-sucrose pairing produces appetitive STM in fed flies. (**H**) OA-VPM3&4 activation during CS-odor presentation, after CS+ odor-sugar pairing, elicits CS-approach. Asterisks denote significant difference, *P* < 0.05. Data presented as mean ± standard error of the mean (S.E.M). Individual data points (circles) correspond to independent biological replicates (8≤*n*≤12). Groups were compared using one-way analysis of variance (ANOVA) with Tukey’s test to correct for multiple comparisons.

## OA-VPM3 activation directs attention to the currently experienced odor

Given OA-VPM3’s considerable connectivity within the MB network, and OA’s documented role in sugar-reinforced appetitive STM (*17–19*), we tested OA-VPM3’s involvement in appetitive STM (Figure. 1D). We used MB022B-GAL4 to express the dominant-negative temperature-sensitive dynamin UAS-*Shibire*^ts1^ (*35–36*) in OA-VPM3 (and OA-VPM4) neurons. Consistent with prior work (*18*), blocking OA-VPM3&4 output by training and testing flies at restrictive 32°C did not affect STM (Figure. 1E), or LTM (Figure. S1B). Blocking only OA-VPM4 (MB113C-GAL4; UAS-*Shi*^ts1^) was also inconsequential for STM (Figure. S1C). Additionally, RNAi-mediated knockdown of *Tβh,* which encodes the OA synthesizing enzyme Tyramine β-hydroxylase (*37*), in OA-VPM3 and OA-VPM4 neurons did not affect STM (Figure. S1D).

We next used an UAS-*dTrpA1* transgene encoding the heat activated cation channel (*38*) to test whether thermogenetic OA-VPM3 activation could implant a memory. Targeted neurons were transiently activated by raising flies to restrictive 32°C. Pairing one of two odors with OA-VPM3&4 activation did not implant an odor preference memory (Figure. 1F&G). However, activating OA-VPM3&4 during the pairing of the second of two odors with sucrose increased the magnitude of appetitive STM formed in hungry and even food-sated flies. Furthermore, OA-VPM3&4 activation with the non-sugar-paired odor (the CS-) formed an appetitive STM for that odor (Figure. 1H), but only if it was preceded by another odor paired with sugar (CS+) and stimulated coincidentally with only one of the two odors (Figure. S2A&B). Activating only OA-VPM4 neurons with odor did not implant a memory or alter sugar-reinforced memory in hungry conditions (Figure. S2C), suggesting the observations with OA-VPM3&4 activation likely result from OA-VPM3. Importantly, OA-VPM3&4 activation during odor-sugar pairing did not produce LTM in starved or sated flies (Figure. S2D&E), nor did activation between training and testing significantly alter memory (Figure. S2F).

Together these data suggest that OA-VPM3 and OA-VPM4 activation temporarily raises the salience of an odor that the flies are currently experiencing. The increased salience can overcome the ordinary hunger-dependence of sugar-reinforced learning and even cause the flies to misassign reward value to an odor that is presented soon after, rather than with, sucrose. Altering salience in a way that specifically impacts STM is consistent with an attentional role of OA-VPM3.

### OAR signaling in MBONs bidirectionally affects appetitive STM

To understand how OA-VPM3 neurons could affect STM, we analyzed their downstream connectivity in the MB network using the Fly-EM hemibrain dataset (*32*). OA-VPM3 neurons preferentially synapse onto Mushroom Body Output Neurons (MBONs) and PPL2 DANs (Table S1, Figure. 2A&B). OA signals through α-type (OAMB, Octα2R), β-type (Octβ1R, Octβ2R, and Octβ3R), and the Oct-TyrR (*16*) receptors (Figure. 2C) which are broadly, but differentially, expressed through the brain. We therefore expressed transgenic RNAi constructs targeting each octopamine receptor (OAR) within 8 relevant MBONs (Figure. 2D) to test whether OA signaling in MBONs influenced memory processing. We did not test the role of PPL2 DANs because a prior study reported no effect on appetitive memory when these neurons were blocked or stimulated during either odor presentation of training (*39*).

**Figure. 2.**
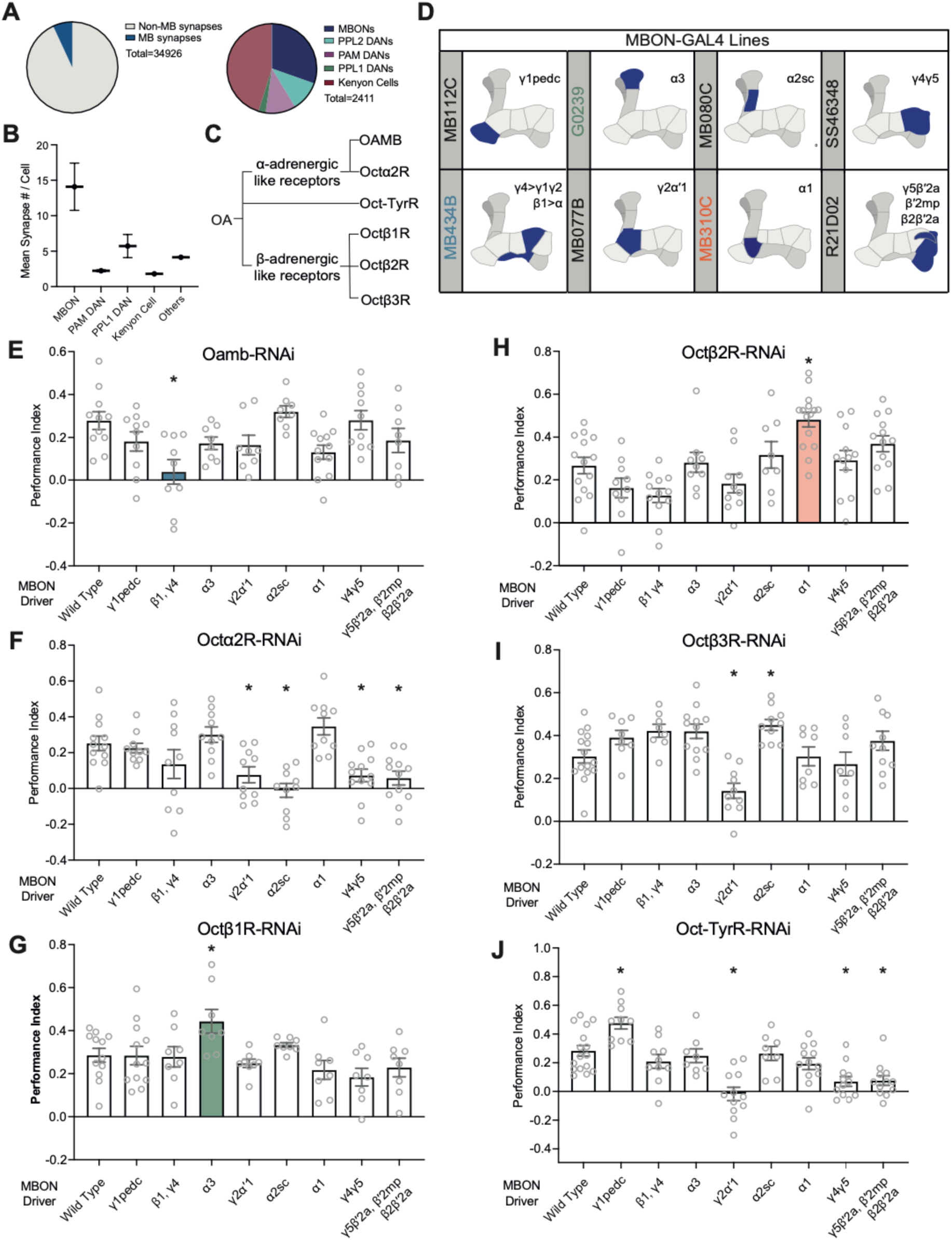
MBON specific OA receptor knock-down bidirectionally modulates appetitive STM. (**A**) Left, 7.4% of putative OA-VPM3 presynapses target Mushroom Body innervating neurons. Right, proportions of the MB-associated synapses made onto different types of MB innervating neurons. (**B**) Mean OA-VPM3 pre-synapses per cell type. (**C**) *Drosophila* OA receptor classification. (**D**) Schematic showing dendritic locations of MBONs screened, and their respective GAL4-drivers. (**E**-**J**) Appetitive STM performance of flies with MBON-GAL4 driven RNAi-mediated knock-down of individual OARs. (**E**&**F**), α-adrenergic-like receptors, OAMB and Octα2R. (**G**-**I**) β-adrenergic-like receptors, Octβ1R, Octβ2R and Octβ3R. (**J**) Oct-TyrR. Asterisks denote significant difference, *P* < 0.05, for groups compared using one-way ANOVA with Dunnett’s test to compare each condition against the heterozygous RNAi control. Data presented as mean ± S.E.M (8≤*n*≤12).

MBON-specific OAR knock-down produced either an increase or a decrease in STM performance. Expressing *Oamb*-RNAi in β1>α/ γ4-MBONs (Figure. 2E, GAL4 controls in Figure. S3A) or *Oct*α*2R*-RNAi in γ2αΩ1, α2sc, γ1pedc, γ5βW2a/β2βW2a MBONs impaired STM (Figure. 2F). In contrast, knocking down *Octβ1R* in MBON-α3 (Figure. 2G) or *Octβ2R* in MBON-α1 (Figure. 2H) enhanced STM. Lastly, two receptors produced opposite effects in different MBONs; *Octβ3R*-RNAi impaired STM when expressed in γ2αΩ1 and enhanced it in α2sc (Figure. 2I) and *Oct-TyrR*-RNAi impaired STM in γ2αΩ1, γ4γ5, y5βW2a/β2βW2a while expression in γ1pedc enhanced STM (Figure. 2J).

Therefore, in 8 of 9 phenotypic cases knock-down of α-receptors in MBONs impaired appetitive STM (Figure. 2E&F) while β-receptor knock-down increased STM in 3 of 4 cases (Figure. 2G-I). Importantly, RNAi of these memory-sensitive receptors driven in αβ_core_ KCs (Figure. S3B) or dorsal fan-shaped body (dFB) neurons (Figure. S3C), which are also postsynaptic to OA-VPM3 neurons, did not alter appetitive STM. OA signaling can therefore increase or decrease appetitive STM performance in a receptor and MBON specific manner. The bi-directional changes of STM observed when manipulating OA receptors in MBONs may account for the apparent neutral effect of blocking OA-VPM3 output (Figure. 1E).

### OA signaling exerts temporally restricted effects on a MB microcircuit

Finding a role for OARs in MBON β1>α, MBON α1, and MBON α3 piqued our attention because the latter two MBONs receive feedforward glutamatergic inhibition from MBON β1>α (Figure. 3A, *32*). Moreover, the hemibrain connectome reveals OA-VPM3 neurons to provide the only direct synaptic source of OA to these MBONs (OA-VPM3 synapses are located on the MBONs’ proximal dendrites or axons) (Figure. 3B). We therefore predicted that OA signaling might shunt MBON activity, thereby rerouting network function to bias STM.

**Figure. 3.**
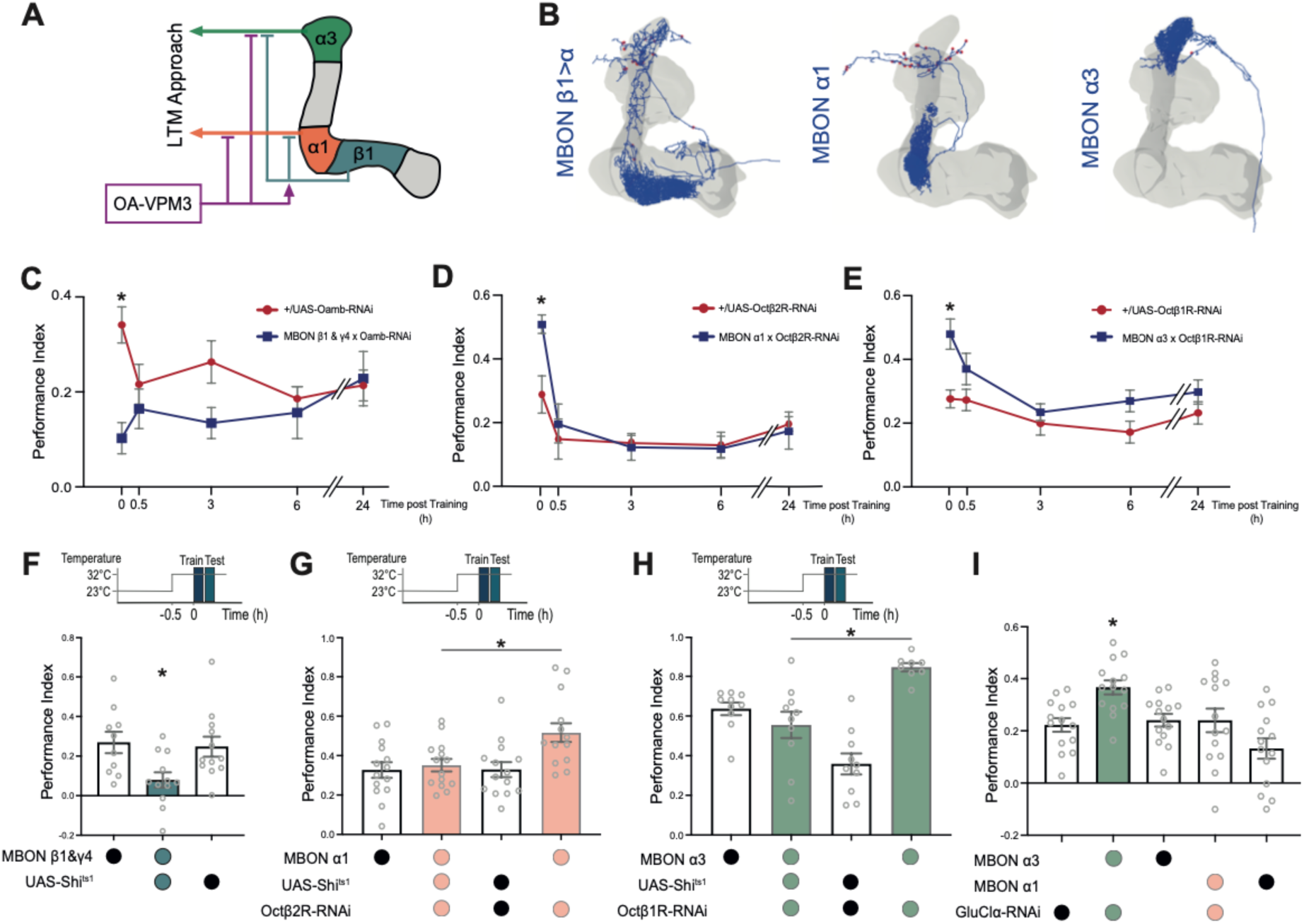
OAR effects in MBONs α1, α3 and β1>α are temporally restricted to STM. (**A**) Diagram of MBON microcircuit downstream of OA-VPM3. (**B**) EM reconstruction of MBON-β1>α, MBON-α1 and MBON-α3 with annotated OA-VPM3 synapses shown in red. (**C**-**E**) Memory timelines of MBON-OAR knockdown flies versus their RNAI controls tested immediately, 0.5, 3, 6 and 24 h after training. (**C**) *Oamb*-RNAi in MBONs β1>α & γ4>γ1>γ2 (MB434B, *n*=8), (**D**) Octβ2R-RNAi in MBON α1 (MB310C, 8≤*n*≤10) and (**E**) Octβ1R-RNAi in MBON α3 (8≤n≤10). Asterisks denote significant difference, *P* < 0.05. Data was compared using a two-way ANOVA or a mixed effect model with Šidák’s multiple comparison test. (**F**-**H**) Top, temperature shifting protocols. (**F**) Blocking MBON β1>α output with UAS-*Shi^t^*^s1^ at 32°C impairs STM. The STM impairment from blocking MBON β1>α output is removed by expressing *Octβ2R*-RNAi in MBON α1 (**G**) or Octβ1R-RNAi in MBON α3 (**H**). (**I**) GluClα RNAi expression in MBON α3, but not MBON α1, enhances STM. Asterisks denote significant difference, *P* < 0.05, groups compared using one-way ANOVA with Tukey’s (**F**), Dunnett’s (**G**&**H**) or Šidák’s (**I**) multiple comparison tests. Data presented as mean ± S.E.M (8≤*n*≤12).

Flies carrying the *Tβh^m18^* mutation are unable to synthesize OA from tyramine (*36*). While *Tβh*^m18^ flies have impaired appetitive STM, their memory from 30 min after training is indistinguishable from that of wild-type flies (*19, 30*). We therefore tested whether knocking-down relevant OARs in the β1>α, α1, and α3 MBONs also produced STM specific effects. *Oamb*-RNAi driven in β1>α and γ4 MBONs exclusively impaired appetitive STM while *Octβ1R*-RNAi in α3 (G0239) or *Octβ2R*-RNA in α1 (MB310C) MBONs enhanced STM (Figure. 3C-E). In each case, memory was indistinguishable from that of their relevant genetic control from 30 min after training, consistent with the STM specificity of OA.

Since MB434B-GAL4 expresses in MBON γ4>γ1γ2 in addition to MBON β1>α, we also expressed *Oamb*-RNAi using a different GAL4 line which labels MBON β1>α (MB433B) and in MBON γ4>γ1γ2 exclusively (VT026001). *Oamb*-RNAi expression in MBON γ4>γ1γ2 did not affect appetitive STM, demonstrating a requirement for OAMB in MBON β1>α for appetitive STM (Figure. S4A).

We next tested whether the effect of knocking-down OA receptors in these 3 MBONs phenocopied MBON inhibition or activation. We again expressed thermogenetic *Shi^ts1^*or *dTrpA1* to respectively inhibit or activate the specific neurons during training and testing. Blocking MBON β1>α output phenocopied *Oamb*-RNAi, indicating that OA normally activates MBON β1>α through OAMB (Figure. 3F).

If the memory enhancement caused by knock-down of Octβ2R in MBON α1 and Octβ1R in MBON α3 results from loss of receptor-dependent neuronal inhibition, then blocking output from these MBONs while the receptors are knocked-down should revert STM performance back to the level of controls. Blocking neural output at restrictive temperature, during training and testing, while driving *Octβ1R*-RNAi and *Octβ2R*-RNAi in MBON α1 and MBON α3 respectively restored control level STM (Figure. 3G & H), while STM remained enhanced in the permissive temperature controls (Figure. S4B & C). Moreover, simply activating MBON α1 during odor-sugar pairing improved STM, phenocopying the effect of *Octβ2R*-RNAi in these neurons (Figure. S4D).

Connectivity suggests that MBON β1>α provides feedforward glutamate-mediated inhibition to MBON α1 and MBON α3 (Figure. 3A; 31). We therefore tested whether this functional connectivity was required for appetitive STM by expressing *GluClα*-RNAi in downstream MBONs (Figure. 4I). As predicted, MBON α3 knock-down of GluClα increased STM, and phenocopied the effect of knocking-down Octβ1R in MBON α3.

**Figure. 4.**
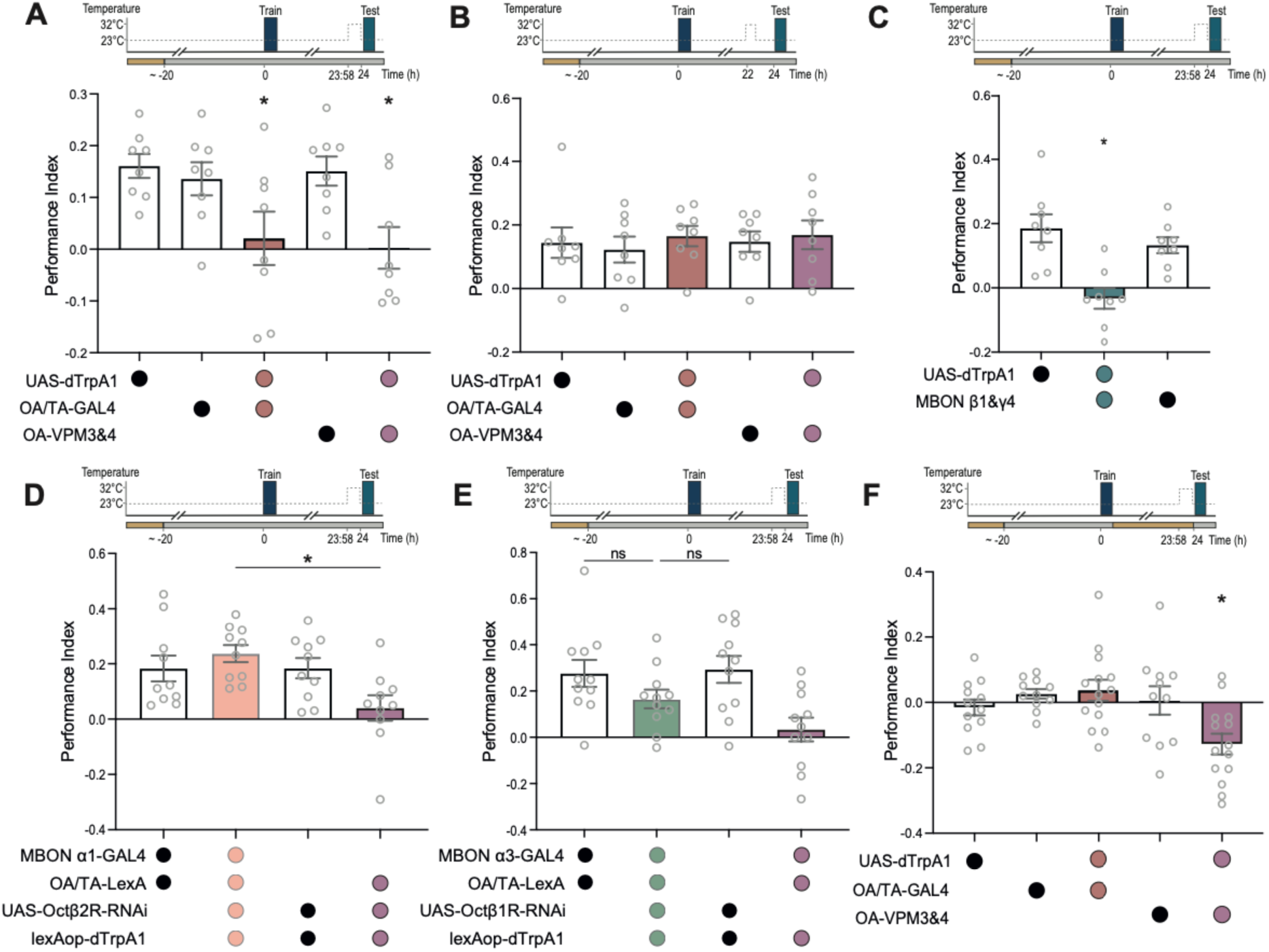
Acute OA-VPM3 activation temporarily inhibits LTM expression. (**A-F**) Top, temperature shifting protocols. (**A**) Appetitive 24 h LTM is abolished in flies where OA/TA (*Tdc2*-GAL4) or OA-VPM3&4 (MB022B) neurons are thermogenetically activated for 2 min prior to testing. (**B**) LTM in flies where OA/TA or OA-VPM3&4 neurons were activated for 2 min, 2 h before testing. (**C**) LTM is abolished in flies in which MBON β1>α (MB434B) are activated for 2 min prior to testing. (**D**&**E**) LTM performance is restored in flies with OA/TA-LexA neurons (*Tdc2*-LexA) artificially activated 2 min before testing if (**D**) *Octβ2R*-RNAi is simultaneously expressed in MBON α1, or (**E**) *Octβ1R*-RNAi in MBON α3. (**F**) 24 h LTM in flies that were fed *ad libitum* after training and received acute OA/TA or OA-VPM3&4 neuron activation for 2 min before testing. Asterisks denote significance, *P* < 0.05, for groups compared using one-way ANOVA with Šidák’s (**A, B, F**), Tukey’s (**C**), or Dunnett’s (**D, E**) multiple comparison test. Data are presented as mean ± S.E.M (8 ≤ n ≤ 14).

Together these results indicate that OA inhibits MBON α1 and MBON α3 via Octβ2R and Octβ1R, respectively. The entopic inhibition of MBONs α1 and α3 during STM is presumably why loss-of-function studies found no STM effects (Figure. S4E&F) and why others concluded that these MBONs have LTM specific roles (*40, 41*).

### OA-VPM3 activity inhibits LTM expression

The α1 and α3 MBONs have documented roles in appetitive LTM retrieval (*40, 41*). We therefore tested whether *dTrpA1*-mediated OA-VPM3 activation inhibited LTM expression. Flies underwent appetitive training and for 2 min prior to testing 24 h LTM, OA-VPM3 neurons were activated by shifting flies to restrictive 32°C. Transient OA-VPM3 and OA-VPM4 (or most OA and TA neurons; *Tdc2*-GAL4) activation inhibited LTM expression (Figure. 4A). However, activating only OA-VPM4 did not alter LTM (Figure. S5).

Importantly, the effect of OA-VPM3 stimulation on LTM expression was transient as LTM performance returned to normal levels 2 h after OA-VPM3 stimulation (Figure. 4B). We therefore concluded that OA-VPM3 activation temporarily ‘masks’ LTM expression. Consequently, transiently activating MBON β1>α also inhibited LTM expression (Figure. 4C).

To more tightly link the LTM masking effect of OA neuron activation with modulation of downstream MBONs, we simultaneously manipulated both classes of neurons.

The broad OA neuron expressing OA/TA-LexA was used to drive LexAop-*dTrpA1* in the same flies expressing OAR RNAi under control of the appropriate MBON-GAL4s. Whereas transient Tdc2 neuron stimulation masked LTM, memory was regained if *Octβ2R*-RNAi was concurrently driven in MBON α1 (Figure. 4D). *Octβ1R*-RNAi in MBON α3 partially recovered LTM following OA/TA neuron activation - the experimental group was not significantly different from the genetic controls or the positive control (Figure. 4E).

We also tested whether transient OA-VPM3 activation affected LTM performance in food-sated flies. We again appetitively trained hungry flies but this time, housed them in food vials between training and testing. Surprisingly, transient activation of only OA-VPM3 and OA-VPM4 neurons, but not all *Tdc2*-GAL4 neurons, led to an inversion of LTM performance, where flies avoided the previously sugar-rewarded odor (Figure. 4F). Such valence inversion suggests that OA-VPM3 may amplify the lack of the sated fly’s desire to seek the sugar-predictive odor.

### OA-VPM3 responds to sucrose and odor presentation

Given artificial activation of OA-VPM3 inhibits LTM, we tested whether the sensory stimuli used during training naturally induce OA-VPM3 activity. We expressed a UAS transgene for the calcium-sensitive fluorescent reporter GCaMP6s (*42*) in OA-VPM3 neurons and recorded calcium transients in OA-VPM3 axons in the superior-medial and -lateral protocerebrum (Figure. 5A). Ingestion of 1M sucrose solution in a 20 s presentation evoked an elevation of intracellular Ca^2+^ in starved flies and a smaller signal in sated flies (Figure. 5B-D). OA-VPM3 responses were not observed when solution only touched the fly’s proboscis or tarsae (data not shown). Moreover, repeated feeding bouts revealed a linear decrease in OA-VPM3’s Ca^2+^ response, consistent with the flies’ diminishing hunger (Figure. 5D).

**Figure. 5.**
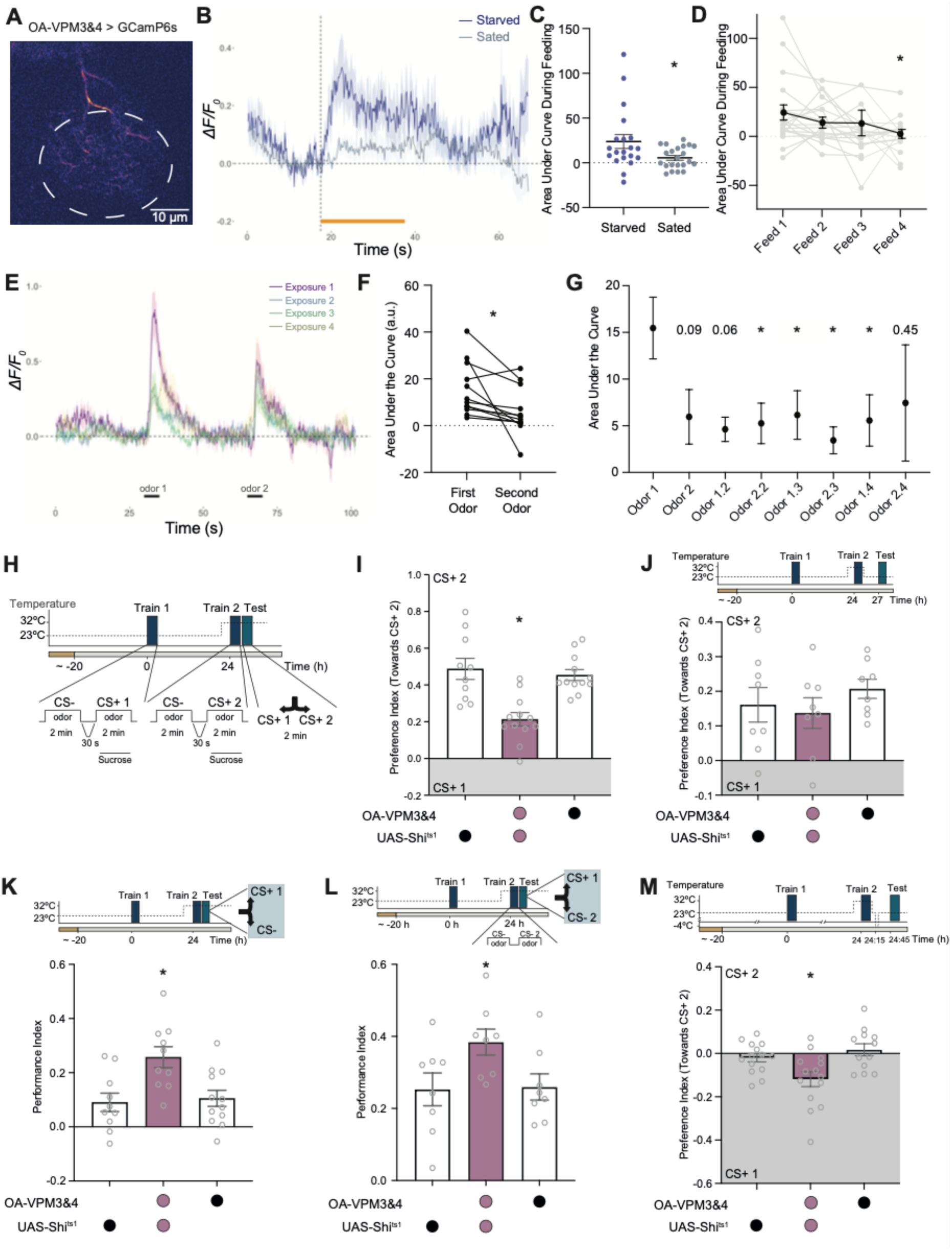
OA-VPM3 prioritizes recent STM by inhibiting expression of remote LTM. (**A**) Representative focal plane for GCaMP6s recording of OA-VPM3 activity. Dashed line encircles presynaptic terminals in quantified region of interest (ROI). (**B**&**C**) Mean sucrose feeding-evoked calcium signal in OA-VPM3 in **B**. sated and starved flies. Orange bar (feeding bout) (**C**) Quantification of area under the curve of OA-VPM3 calcium responses during the feeding bout. Data compared with Mann-Whitney test. (**D**) Calcium responses diminish over successive feeding bouts in starved flies. Asterisk denotes significance, *P*<0.05 when implementing a mixed-effects analysis with Dunnett’s test to compare responses to Feed 1 (8≤*n*≤12). (**E**) Mean odor-evoked calcium responses in OA-VPM3 in sated flies. Each trial consisted of 2 odor presentations and each flies received 4 successive trials with a ∼2 min inter-trial interval. (**F**) Calcium response of the first odor presentation compared to the next presentation using a paired t-test (**G**) Mean area under the curve for successive odor presentations across exposure trials. Data presented as mean ± S.E.M and difference tested using a mixed-effect model with Dunnett’s test to compare to odor 1 response. (**H**) Protocol to test between remote LTM (trained CS+1 v CS-) acquired 24 h prior to testing versus a recent STM (trained CS+2 v CS-) acquired immediately before testing. Flies were tested for preference between CS+2 and CS+1, or for only remote CS+1 v CS-Temperature was raised to restrictive 32°C at particular times to inactivate neurons expressing UAS-*Shi*^ts1^. (**I**-**M**) Top, train, test and temperature shifting protocols. (**I**) Preference for recent CS+2 over remote CS+1 is compromised by blocking OA-VPM3&4 during CS+2 training and memory preference testing. (**J**) Preference for recent CS+2 over CS+1 is not impaired when OA-VPM3&4 are blocked during CS+2 training but CS+2 v CS+1 is tested 3 h later (3h CS+1 memory vs 27 h CS+2 memory). (**K**) Blocking OA-VPM3&4 during CS+2 training enhances expression of remote CS+1 LTM (test CS+1 v CS-). (**L**) Blocking OA-VPM3&4 during a recent ‘mock’ training (odors only, CS- and CS-2) enhances expression of remote LTM (test CS+1 v CS-2). (**M**) Combining OA-VPM3&4 block during CS+2 training and cold-shock anesthesia 15 min after CS+2 training reveals approach to CS+1 LTM when flies are tested 30 min later (CS+ 1 v CS+ 2). Asterisks denote significant difference, *P* < 0.05, for groups compared using one-way ANOVA with Tukey’s test. Data are mean ± S.E.M (8≤*n*≤14).

We next tested whether OA-VPM3 respond to odors. Fed flies received two 5 s pulses of 2 different odors. This regimen was repeated 4 times per fly (Figure. 5E). OA-VPM3 responded more strongly to odor presentation than to sucrose feeding. However, the 6 subsequent odor-evoked responses to odor were of significantly reduced amplitude to that of the first exposure, demonstrating that VPM3 undergoes rapid odor-general adaptation (Figure. 5F&G). Reduced responding was time limited as the magnitude of the 7^th^ odor response had returned to that of the first presentation. (Figure. 5G, Figure. S6A-D). Therefore, OA-VPM3 neurons show state-modulated responses to sucrose whereas their odor responses adapt with a refresh rate of ∼15 min. Such a response profile may allow OA-VPM3-released OA to set an attentional window.

### OA-VPM3 activity promotes expression of recent over remote memories

Knowing that OA-VPM3 activation promotes STM and inhibits LTM, and that it is naturally activated by the stimuli used in training, led us to revisit the lack of a STM phenotype when blocking OA-VPM3 neurons (Fig 1.E). We reasoned that OA-VPM3 activity during learning could preferentially promote expression of new memories (STM) by simultaneously suppressing use of older (LTM) food-seeking memories.

We tested for prioritization of STM using a consecutive training protocol (Figure. 5H). Flies were first trained by presenting one odor with nothing (CS-) for 2 min, then pairing odor 1 with sugar (CS+1). 24 h later the same flies were trained again with one odor with nothing (CS-, same odor as in first training) then a different odor with sugar (CS+2). When subsequently tested between ‘remote LTM’ (CS+1) odor and ‘recent STM’ (CS+2) odor wild-type flies showed clear preference for CS+2 (Figure. 5I). However, blocking OA-VPM3 output with *Shi*^ts1^ during the second training session impaired preference for recent CS+2 over remote CS+1 (Figure. 5I; OA-VPM4 block was inconsequential Figure. S7A). If memory was instead tested 3 h after CS+2 training the OA-VPM3 effect had abated – demonstrating that OA-VPM3’s contribution to preferring recent CS+2 memory is time-limited (Figure. 5J).

Blocking OA-VPM3 during the second training session also enhanced remote CS+1 memory if performance was tested between remote CS+1 and the CS-odor (Figure. 5K). Led by our observation that OA-VPM3 is odor activated, we withheld sucrose from the second training session and instead only presented the two odors (CS-1 and CS-2). Blocking OA-VPM3 after these ‘mock training’ odor exposures also enhanced the flies’ expression of remote memory (CS+1, when tested between CS+1 and CS-2) (Figure. 5L). In addition, when flies were tested 24 h after the second training session no significant preference was apparent between remote CS+2 and remote CS+1, indicating that two remote memories (one 24 h and other 48 h) have equivalent predictive value (Figure. S7B). Further, only sugar presentation prior to testing (Fig S7C) or blocking OA-VPM3 during the first training session had no effect on memory (Figure. S7D), and the recently acquired STM after consecutive training was also unaffected by blocking OA-VPM3 (Figure. S7E).

Appetitive STM is labile and susceptible to cold-shock anesthesia, while appetitive LTM is cold-shock resistant and therefore operationally ‘consolidated’ (*43, 44*). We therefore used anesthesia after training to further test the hypothesis that OA-VPM3 removes interference from LTM to promote STM (Figure. 5M). We again trained flies over two days with two consecutive sessions and blocked OA-VPM3 output during the second training. Flies were then briefly cold-shock anesthetized 15 min after the second training, then given 30 min to recover before testing preference between remote CS+1 LTM and recent (∼45 min) CS+2 STM. In both the genetic controls, memory performance was indistinguishable from 0, consistent with CS+1 LTM and 45 min ∼CS+2 STM being in balance. However, CS+1 LTM emerged in flies in which OA-VPM3 neurons had been blocked during the second training, presumably because CS+2 STM was absent due to cold-shock treatment, while LTM was unmasked by removing OA-VPM3-mediated suppression. Together these results suggest that OA-VPM3 activity temporarily reconfigures the MBON network to favor STM expression by simultaneously masking LTM expression.

## Discussion

In this study we demonstrate an attention-like role for a specific pair of broadly projecting OA neurons (OA-VPM3). Both their artificial and odor (and learning) evoked activity promotes expression of new experience/memory in favor of remote experience. We localized key sites of OA action by removing different OARs from pairs of MBONs that are directly postsynaptic to OA-VPM3 neurons. MBON identity and the known signaling modes of the respective OARs they employ led us to determine that OA-VPM3 activity facilitates MBON pathways for expression of STM (via α-type receptors) while simultaneously inhibiting those directing LTM (via β-type receptors). To our knowledge, finding such a STM promoting mechanism for OA provides the first clear example of an attentional ‘network reset’ function proposed for the analogous mammalian noradrenergic system (*2, 45*).

Stimulus-evoked responses of OA-VPM3 are transient but the released OA will produce different temporal effects in each target location as determined by the OAR pathways it engages there. Importantly, our behavior experiments show that the STM promoting/LTM inhibiting effects are transient and reversible indicating OAR activation masks MB output without affecting DAN induced plasticity at KC>MBON junctions. It will be interesting to determine whether the OA-directed attentional process is and/or can be modality selective, and how attention like processes integrate across timescales (*46*).

Several studies have reported hunger-dependent peptidergic and monoaminergic modulation of the MB network to be key to promote innate and learned appetitive behaviors (*24, 47–49*). We propose that the temporary OA-directed network reset functions on top of these other longer duration mechanisms to permit the fly to attend to their most recently acquired odor-food association to optimize active foraging. In this way, behavioral states track featural information, such as timing of acquisition, to bias the fly’s preference and break indecision between otherwise equivalent resource-seeking memories.

## Acknowledgments

We thank Nils Otto for help with connectomics code and members of the Waddell lab for discussion. We thank the Vienna Drosophila Resource Center and the Bloomington Stock Center for flies. **Funding:** I.K was funded by the Biotechnology and Biological Sciences Research Council (DDT00230). S.W. was funded by a Wellcome Principal Research Fellowship (200846), a Wellcome Discovery Award (225192), an ERC Advanced Grant (789274), and Wellcome Collaborative Awards (203261 and 209235).

## Author contributions

Designed research I.K., S.W., Performed research I.K., Analyzed data I.K., S.W., Resources S.W. Writing I.K., S.W., Supervision S.W., Funding Acquisition S.W.

## Data and materials availability

All data are available in the manuscript or in the supplementary materials. Code and raw data are available on request.

## Materials and Methods

### Fly strains

*Drosophila melanogaster* strains used in this study were maintained on standard cornmeal-agar food under a 12:12 h light: dark cycle. Strains were reared with 40-50% relative humidity and, other than crosses expressing thermosensitive transgenes, raised at 25 °C. Crosses expressing UAS-*dTrpA1* were grown at 18 °C and those expressing UAS-*Shi*^ts1^ at 23 °C. Mixed sex populations and individuals were used for behavior and imaging experiments, respectively. Canton-S (CS) flies were used as the wild-type strain. The following transgenic strains were used, in order of appearance in the figures.

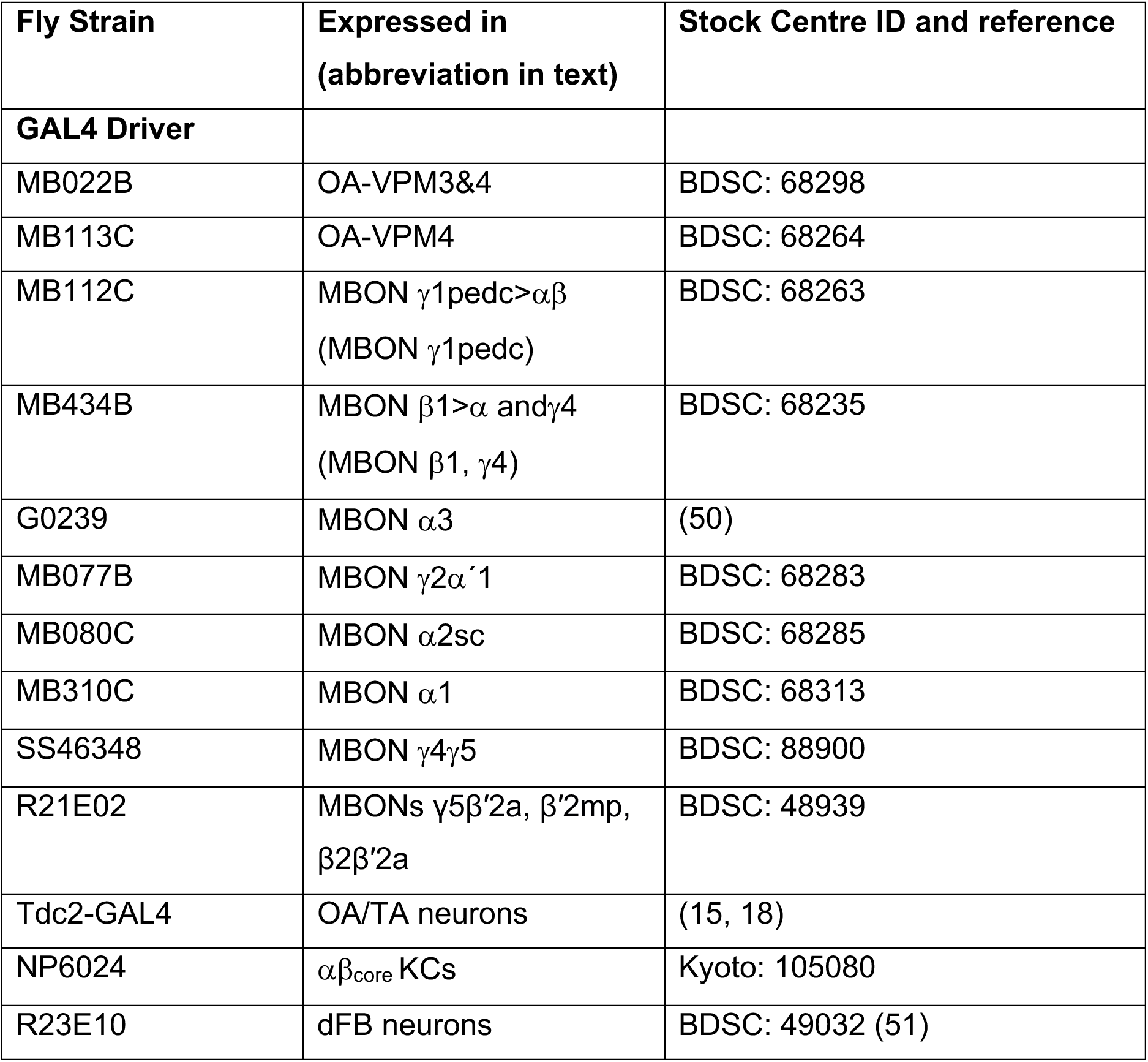
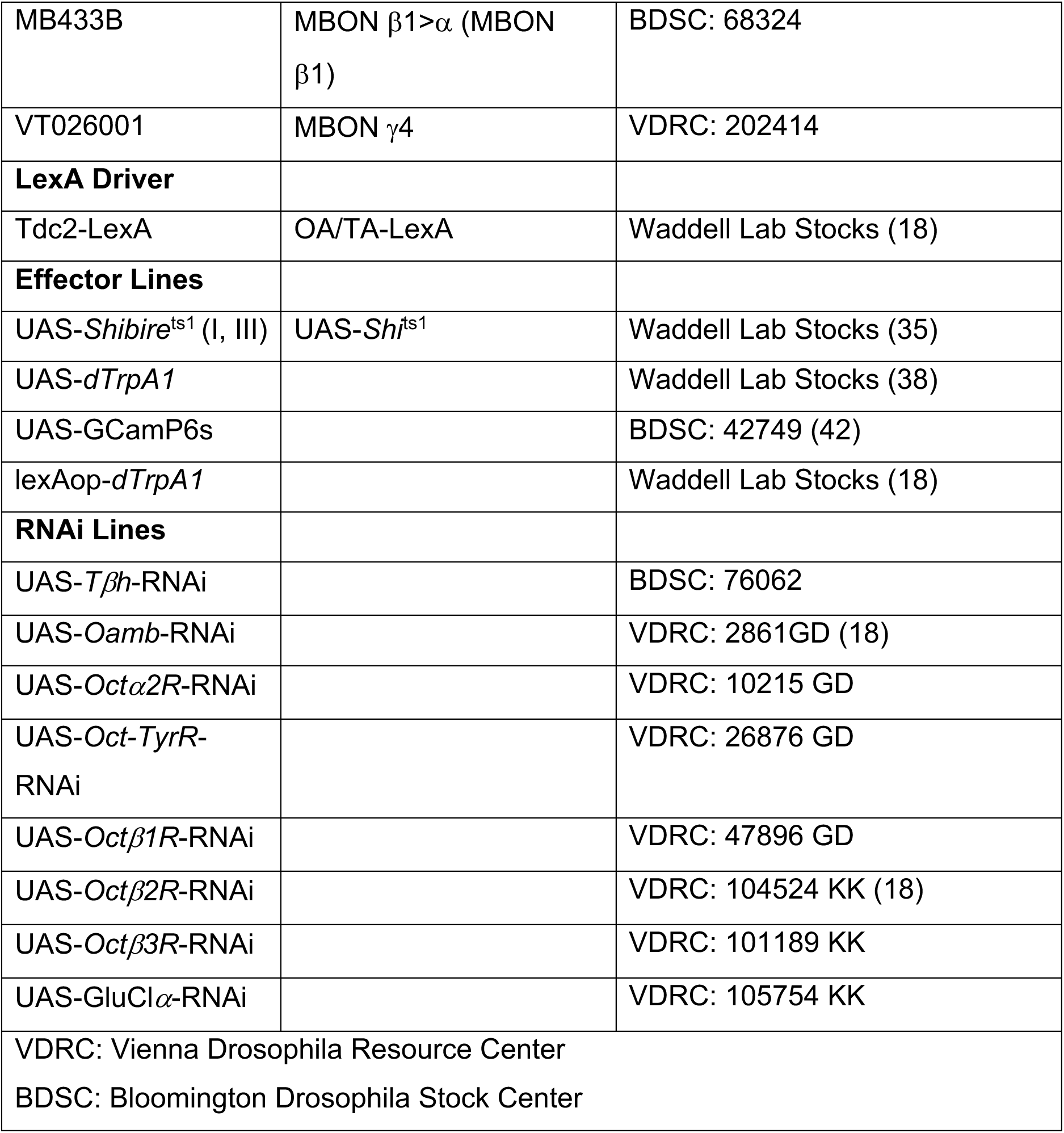

## Connectomics Analysis

The relevant connectomes and associated information were downloaded from either neuPrint+ (hemibrain v1.2.1) or Codex FlyWire (fafb v630). Synaptic connectivity information and synapse locations were downloaded from neuprintr (hemibrain v1.2.1) and analyzed and indexed using a custom Python and R script. Sankey diagram and mesh diagrams of relevant neurons were generated using adapted R scripts.

## Behavior

For experimental crosses, 5-8 male flies from the specified GAL4 line were crossed to 20-25 virgin females from the UAS lines, with the exception of Figures 3G, 3H, 4D, and 4E, where GAL4 females were crossed to effector males. For heterozygous parental controls, GAL4 or UAS lines were crossed to wild-type CS females or males, consistent with the crosses in the experimental groups.

Behavioral T-maze experiments were conducted using adapted versions of previously described protocols (47). 60-100 group-housed, mixed sex flies that were 2-9 days old were used per trial. Unless specified, experiments were conducted at 23 °C and 50-70% relative humidity. For thermogenetic manipulations with either UAS-*dTrpA1* or UAS-*Shi*^ts1^, temperature was raised to the restrictive 31-33°C at the times specified in the provided schematic protocols. Odors used were 3-octanol (OCT, 1:10^3^), 4-methylcyclohexanol (MCH, 1:10^3^), and isoamyl acetate (IAA, 2:10^3^), diluted in mineral oil in the concentrations specified in parentheses. Odors were bubbled at a consistent flow rate by adjusting vacuum pressure through the T-maze prior to the experiment.

Unless specified, flies were starved for 18-24 h prior to appetitive conditioning. Approximately 80-100 flies were aliquoted in 25 ml vials containing 2 ml pellet of 1% agar to keep the flies hydrated. A 20 x 60 mm strip of Whatman filter paper was added to each vial. Flies were kept food deprived throughout the entire experiment unless otherwise specified. In the cases where flies were fed, ∼80-100 flies were placed in a 25 ml vial with standard food and a 20 x 60 mm strip of Whatman filter paper.

For single trial appetitive conditioning, starved flies were exposed to a CS-odor for two min in a tube lined with Whatman filter paper that had been previously soaked with tap water and subsequently dried. Flies were allowed to rest for 30 s post CS-exposure in the same tube with clean air. Flies were then moved to a new tube lined with Whatman filter paper coated in dried sucrose (paper dipped in saturated 5.8M sucrose solution then dried) and exposed to a CS+ odor for 2 min. For UAS-*Shi*^ts1^ experiments, flies were moved to restrictive 32-33 °C temperature 30 min prior to targeted time to allow them to acclimatize to the new temperature. UAS-*dTrpA1* flies were moved to permissive temperatures of 31-32 °C at specified times for acute neuronal activation.

Memory performance was measured by testing for odor preference between 2 odors in the dark. In most experiments, Performance Index (PI) was calculated as the weighted difference between flies that picked the CS+ odor over the CS-odor using the formula:

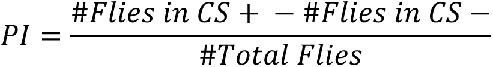

The PI score across two independent, reciprocally trained trials (either odor as CS+) were averaged as one *n* and are each represented by a point on the respective graphs. In the case where multiple odors were used, tested odors are indicated in the protocol schematic and PI score is calculated with positive values towards either the previously reinforced odor or the more recent odor in cases where both odors were paired with sucrose reward. In all experiments the two testing odors were OCT and MCH.

## In-vivo calcium imaging

Group housed flies were raised at 25 °C and 2–8-day old flies, of either sex, were used. Imaging experiments were conducted as previously described (52). Flies were briefly anesthetized on ice and mounted on a 3D printed platform using UV glue (Bondic) such that the proboscis, antennae, and legs could move freely. The head capsule was opened under 1 ml sugar-free HL3-like solution (140 mM NaCl, 2 mM KCl, 4.5 mM MgCl_2_, 1.5 mM CaCl_2_, 5 mM HEPES-NaOH, pH 7.1, 275 mOsm kg^−1^). Post-surgery, the buffer was removed and a droplet of ∼10 μl of 1% agarose dissolved in sugar-free saline was placed over the exposed brain to help minimize motion while allowing the proboscis to move. The chamber was filled with 1 ml sugar-free saline and placed under a two-photon microscope.

For feeding experiments, a 10 μl droplet of 1M sucrose solution with 0.4% brilliant blue dye on a 3D printed tee was manually raised using a micromanipulator to the fly’s proboscis (53, 54). The time of feeding was manually noted, and a feeding bout was ended 20 s after the start of feeding. Flies were scored for blue dye in their abdomen and included in the analysis if they had ingested the sucrose solution. For odor delivery, a custom-built odor delivery system was used as previously described (53, 56). Two pulses of odor (OCT and MCH) 30 s apart were switched into a clear air stream. Each pulse of odor lasted 5 s and the order in which odors were presented was swapped between flies but kept consistent for any given fly.

A customized Scientifica two-photon microscope was equipped with a 40× 0.8 NA water-immersion objective (Olympus) and a dichroic beamsplitter (BrightLine, Semrock) with a green (500/15 nm, Semrock) filter followed by a GaAsP PMT (Hamamatsu) to detect GCaMP6s signal. A Ti:sapphire laser (Chameleon Ultra II, Coherent) centered on 910 nm was used to excite fluorescence using ∼140 fs pulses at 80 MHz repetition rate. Images of 256 × 256 pixels were acquired at 5.95 Hz under the control of ScanImage 3.8 software via MATLAB (MathWorks, release 2012a).

Images were processed and manually segmented using FIJI where ROIs were manually selected to calculate summed fluorescence and background subtracted for that frame. As there was minimal drift over the course of imaging, brains were not registered to a template. Fluorescence traces are depicted as ΔF/F_0_, where the baseline or F_0_ was calculated as the mean signal collected over 5 s prior to stimulus exposure. Area under the curve was calculated as the difference from baseline during stimulus exposure. Graphs, area under the curve, and max peak heights were calculated on R 4.3.2.

### Statistics

Statistical tests were implemented with GraphPad Prism 9.4.1. All behavioral data was analyzed with an unpaired t-test, ordinary one-way ANOVA or two-way ANOVA/mixed-effects model with a Tukey (all groups compared), Šidák (select groups compared), or Dunnett’s (groups compared to a single control) test to correct for multiple comparisons (see table below). The post-hoc test was selected based on the pairwise comparisons being made.

For behavior experiments, biological replicates are represented as single points and each replicate (*n*) is the average between two reciprocally trained groups. Therefore, the population behavior of ∼ 120-200 flies accounts for each *n*. For calcium imaging experiments, each *n* represents a single fly. Paired t-test or repeated measures tests were used to analyze repeat feeding and odor exposures in the same fly. All experiments were repeated across at least three separate sessions on different days.

The following statistical tests were used to report significance in the relevant figures:

| Figure | Test and correction/<br>multiple comparison test | Pairwise Comparison | P-value |
| --- | --- | --- | --- |
| **Figure 1** |  |  |  |
| 1E | One-way ANOVA<br>- Tukey post-hoc | VPM3&4 x *Shi<sup>ts1</sup>* vs VPM3&4<br>VPM3&4 x *Shi<sup>ts1</sup>* vs *Shi<sup>ts1</sup>* | 0.5190<br>0.9853 |
| 1F | One-way ANOVA<br>- Tukey post-hoc | VPM3&4 x *dTrpA1* vs VPM3&4<br>VPM3&4 x *dTrpA1* vs *dTrpA1*<br>VPM3&4 x *dTrpA1* vs VPM3&4 x *dTrpA1* (no sucrose) | 0.0285<br>0.0003<br><0.0001 |
| 1G | One-way ANOVA<br>- Tukey post-hoc | VPM3&4 x *dTrpA1* vs VPM3&4<br>VPM3&4 x *dTrpA1* vs *dTrpA1*<br>VPM3&4 x *dTrpA1* vs VPM3&4 x *dTrpA1* (no sucrose) | 0.0005<br>0.0018<br>0.0429 |
| 1H | One-way ANOVA<br>- Tukey post-hoc | VPM3&4 x *dTrpA1* vs VPM3&4<br>VPM3&4 x *dTrpA1* vs *dTrpA1* | 0.0038<br>0.0394 |
| **FigureS1** |  |  |  |
| S1B | One-way ANOVA<br>- Tukey post-hoc | VPM4 x *Shi<sup>ts1</sup>* vs VPM4<br>VPM4 x *Shi<sup>ts1</sup>* vs *Shi<sup>ts1</sup>* | 0.0949<br>0.9712 |
| S1C | One-way<br>ANOVA<br>- Tukey<br>post-hoc | VPM3&4 x Tβh-RNAi vs VPM3&4<br>VPM3&4 x Tβh-RNAi vs Tβh-RNAi | 0.9637<br>0.9420 |
| S1D | One-way<br>ANOVA<br>Tukey post-hoc | VPM3&4 x *Shi*<sup>ts1</sup> vs VPM3&4<br>VPM3&4 x *Shi*<sup>ts1</sup> vs *Shi*<sup>ts1</sup> | 0.2094<br>0.6732 |
| **FigureS2** |  |  |  |
| S2A | One-way<br>ANOVA<br>- Tukey<br>post-hoc | VPM4 x *dTrpA1* (no sucrose) vs<br>VPM4<br>VPM4 x *dTrpA1* (no sucrose) vs<br>*dTrpA1*<br>VPM4 x *dTrpA1* (no sucrose) vs<br>VPM4 x *dTrpA1* | 0.0216<br>0.0258<br>0.0091 |
| S2B | One-way<br>ANOVA<br>- Tukey<br>post-hoc | VPM3&4 x *dTrpA1* vs VPM3&4<br>VPM3&4 x *dTrpA1* vs *dTrpA1* | 0.2161<br>0.3886 |
| S2C | One-way<br>ANOVA<br>- Tukey<br>post-hoc | VPM3&4 x *dTrpA1* vs VPM3&4<br>VPM3&4 x *dTrpA1* vs *dTrpA1* | 0.3698<br>0.9130 |
| S2D | One-way<br>ANOVA<br>- Tukey<br>post-hoc | VPM3&4 x *dTrpA1* vs VPM3&4<br>VPM3&4 x *dTrpA1* vs *dTrpA1*<br>VPM3&4 x *dTrpA1* vs VPM3&4 x<br>*dTrpA1* (no sucrose) | 0.8732<br>0.7300<br>0.3811 |
| S2E | One-way<br>ANOVA<br>- Tukey<br>post-hoc | VPM3&4 x *dTrpA1* (no sucrose)<br>vs VPM3&4<br>VPM3&4 x *dTrpA1* (no sucrose)<br>vs *dTrpA1*<br>VPM3&4 x *dTrpA1* (no sucrose)<br>vs VPM3&4 x *dTrpA1* | 0.0006<br>0.0048<br>0.0007 |
| **Figure 2** |  |  |  |
| 2E | One-way ANOVA<br>- Dunnett's test | MBON $\beta 1, \gamma 4$ vs Wild Type | 0.001 |
| 2F | One-way ANOVA<br>- Dunnett's test | MBONs $\gamma 5\beta'2a, \beta'2mp, \beta 2\beta'2a$ vs Wild Type<br>MBON $\gamma 2\alpha'1$ vs Wild Type<br>MBON $\alpha 2sc$ vs Wild Type<br>MBON $\gamma 1\gamma 2$ vs Wild Type | 0.0128<br>0.0437<br>0.0006<br>0.0259 |
| 2G | One-way ANOVA<br>- Dunnett's test | MBON $\gamma 1pedc$ vs Wild Type<br>MBONs $\gamma 5\beta'2a, \beta'2mp, \beta 2\beta'2a$ vs Wild Type<br>MBON $\gamma 2\alpha'1$ vs Wild Type<br>MBON $\gamma 1\gamma 2$ vs Wild Type | 0.0064<br>0.0013<br><0.0001<br>0.0009 |
| 2H | One-way ANOVA<br>- Dunnett's test | MBON $\alpha 3$ vs Wild Type | 0.03773 |
| 2I | One-way ANOVA<br>- Dunnett's test | MBON $\alpha 1$ vs Wild Type | 0.0007 |
| 2J | One-way ANOVA<br>- Dunnett's test | MBON $\gamma 2\alpha'1$ vs Wild Type<br>MBON $\alpha 2sc$ vs Wild Type | 0.0084<br>0.019 |
| **Figure 3** |  |  |  |
| 3C | Two-way repeated measures ANOVA with Geisser- | MBON $\beta 1, \gamma 4$ x *Oamb*-RNAi vs *Oamb*-RNAi<br>- Immediate Memory (0 h) | 0.0016 |
|  | Greenhouse<br>Correction<br>- Šidák |  |  |
| 3D | Two-way<br>repeated<br>measures<br>ANOVA with<br>Geisser-Greenhouse<br>Correction<br>- Šidák | MBON $\alpha 1$ x *Oct$\beta 2R$-RNAi* vs<br>*Oct$\beta 2R$-RNAi*<br>- Immediate Memory (0 h) | 0.0072 |
| 3E | Mixed-effect<br>model with the<br>Geisser-Greenhouse<br>Correction<br>- Šidák | MBON $\alpha 3$ x *Oct$\beta 1R$-RNAi* vs<br>*Oct$\beta 1R$-RNAi*<br>- Immediate Memory (0 h) | 0.0167 |
| 3F | One-way<br>ANOVA<br>- Tukey<br>post-hoc | MBON $\beta 1, \gamma 4$ x *Shi<sup>ts1</sup>* vs MBON<br>$\beta 1, \gamma 4$<br>MBON $\beta 1, \gamma 4$ x *Shi<sup>ts1</sup>* vs *Shi<sup>ts1</sup>* | 0.0248<br>0.0387 |
| 3G | One-way<br>ANOVA<br>- Dunnett's<br>test | MBON $\alpha 1$ x *Oct$\beta 2R$-RNAi* vs<br>MBON $\alpha 1$<br>MBON $\alpha 1$ x *Oct$\beta 2R$-RNAi* vs<br>*Oct$\beta 2R$-RNAi*; *Shi<sup>ts1</sup>*<br>MBON $\alpha 1$ x *Oct$\beta 2R$-RNAi* vs<br>MBON $\alpha 1$ x *Oct$\beta 2R$-RNAi*; *Shi<sup>ts1</sup>* | 0.0038<br>0.0042<br>0.0131 |
| 3H | One-way<br>ANOVA<br>- Dunnett's<br>test | MBON $\alpha 3$ x *Oct$\beta 1R$-RNAi* vs<br>MBON $\alpha 3$<br>MBON $\alpha 3$ x *Oct$\beta 1R$-RNAi* vs<br>*Oct$\beta 1R$-RNAi*; *Shi<sup>ts1</sup>* | 0.0154<br><0.0001<br>0.0007 |
| | | MBON $\alpha 3$ x *Oct$\beta 1R$-RNAi* vs<br>MBON $\alpha 3$ x *Oct$\beta 1R$-RNAi; Shi<sup>ts1</sup>* | |
| 3I | One-way<br>ANOVA<br>- Šidák | MBON $\alpha 3$ x *GluClα-RNAi* vs<br>MBON $\alpha 3$<br>MBON $\alpha 3$ x *GluClα-RNAi* vs<br>*GluClα-RNAi* | 0.0378<br>0.0130 |
| **Figure S4** |  |  |  |
| S4A | One-way<br>ANOVA<br>- Šidák | MBON $\beta 1$ x *Oamb-RNAi* vs<br>*Oamb-RNAi*<br>MBON $\beta 1$ x *Oamb-RNAi* vs<br>MBON $\beta 1$<br>MBON $\gamma 4$ x *Oamb-RNAi* vs<br>*Oamb-RNAi*<br>MBON $\gamma 4$ x *Oamb-RNAi* vs<br>MBON $\gamma 4$ | <0.0001<br><0.0001<br>0.0095<br>0.9985 |
| S4B | One-way<br>ANOVA<br>- Dunnett's<br>test | MBON $\alpha 1$ x *Oct$\beta 2R$-RNAi* vs<br>MBON $\alpha 1$<br>MBON $\alpha 1$ x *Oct$\beta 2R$-RNAi* vs<br>*Oct$\beta 2R$-RNAi; Shi<sup>ts1</sup>*<br>MBON $\alpha 1$ x *Oct$\beta 2R$-RNAi* vs<br>MBON $\alpha 1$ x *Oct$\beta 2R$-RNAi; Shi<sup>ts1</sup>* | 0.0086<br>0.0003<br>0.7032 |
| S4C | One-way<br>ANOVA<br>- Dunnett's<br>test | MBON $\alpha 3$ x *Oct$\beta 1R$-RNAi* vs<br>MBON $\alpha 1$<br>MBON $\alpha 3$ x *Oct$\beta 1R$-RNAi* vs<br>*Oct$\beta 1R$-RNAi; Shi<sup>ts1</sup>*<br>MBON $\alpha 3$ x *Oct$\beta 1R$-RNAi* vs<br>MBON $\alpha 3$ x *Oct$\beta 1R$-RNAi; Shi<sup>ts1</sup>* | 0.0002<br><0.0001<br>0.0841 |
| S4D | One-way<br>ANOVA | MBON $\alpha 1$ x *dTrpA1* vs MBON $\alpha 1$<br>MBON $\alpha 1$ x *dTrpA1* vs *dTrpA1* | 0.0009<br>0.0001<br><0.0001 |
| | - Tukey<br>post-hoc | MBON $\alpha 1$ x *dTrpA1* vs MBON $\alpha 1$<br>x *dTrpA1* (no sucrose) | |
| **Figure 4** |  |  |  |
| 4A | One-way<br>ANOVA<br>- Šidák | OA/TA-GAL4 x *dTrpA1* vs<br>OA/TA-GAL4<br>OA/TA-GAL4 x *dTrpA1* vs<br>*dTrpA1*<br>OA-VPM3&4 x *dTrpA1* vs OA-VPM3&4<br>OA-VPM3&4 x *dTrpA1* vs *dTrpA1* | 0.0175<br>0.0299<br>0.0260<br>0.0158 |
| 4C | One-way<br>ANOVA<br>- Tukey<br>post-hoc | MBON $\beta 1, \gamma 4$ x *dTrpA1* vs MBON<br>$\beta 1, \gamma 4$<br>MBON $\beta 1, \gamma 4$ x *dTrpA1* vs<br>*dTrpA1* | 0.0070<br>0.0005 |
| 4D | One-way<br>ANOVA<br>- Dunnett's<br>Test | MBON $\alpha 1$;OA/TA-LexA x<br>*Octβ2R-RNAi*;LexAop-*dTrpA1* x<br>*Octβ2R-RNAi*;LexAop-*dTrpA1*<br>MBON $\alpha 1$;OA/TA-LexA x<br>*Octβ2R-RNAi*;LexAop-*dTrpA1* x<br>MBON $\alpha 1$;OA/TA-LexA<br>MBON $\alpha 1$;OA/TA-LexA x<br>*Octβ2R-RNAi*;LexAop-*dTrpA1* x<br>OA/TA-LexA x *Octβ2R-*<br>*RNAi*;LexAop-*dTrpA1* | 0.6833<br>0.6723<br>0.0043 |
| 4E | One-way<br>ANOVA<br>- Dunnett's<br>Test | MBON $\alpha 3$;OA/TA-LexA x<br>*Octβ1R-RNAi*;LexAop-*dTrpA1* x<br>*Octβ1R-RNAi*;LexAop-*dTrpA1*<br>MBON $\alpha 3$;OA/TA-LexA x<br>*Octβ1R-RNAi*;LexAop-*dTrpA1* x<br>MBON $\alpha 3$;OA/TA-LexA<br>MBON $\alpha 3$;OA/TA-LexA x<br>*Octβ1R-RNAi*;LexAop-*dTrpA1* x | 0.2178<br>0.3235<br>0.1988 |
|  |  | OA/TA-LexA x *Octβ1R*-RNAi;LexAop-*dTrpA1* |  |
| 4F | One-way ANOVA<br>- Šidák | OA-VPM3&4 x *dTrpA1* vs OA-VPM3&4<br>OA-VPM3&4 x *dTrpA1* vs *dTrpA1* | 0.0143<br>0.0450 |
| **Figure 5** |  |  |  |
| 5C | Mann-Whitney test | Starved vs Fed | 0.0491 |
| 5D | Mixed-effects analysis with Geisser-Greenhouse correction and Dunnett's multiple comparison test | Feed 1 vs Feed 2<br>Feed 1 vs Feed 3<br>Feed 1 vs Feed 4 | 0.7832<br>0.6712<br>0.0308 |
| 5F | Paired t-test | Odor 1 vs Odor 2 | 0.0198 |
| 5G | Mixed-effects analysis with Geisser-Greenhouse correction and Dunnett's multiple comparison test | Odor1 vs. Odor2<br>Odor1 vs. Odor1.2<br>Odor1 vs. Odor2.2<br>Odor1 vs. Odor1.3<br>Odor1 vs. Odor2.3<br>Odor1 vs. Odor1.4<br>Odor1 vs. Odor2.4 | 0.0938<br>0.0656<br>0.0089<br>0.0139<br>0.0116<br>0.0097<br>0.4541 |
| 5I | One-way ANOVA<br>- Tukey post-hoc | VPM3&4 x *Shi*<sup>ts1</sup> vs VPM3&4<br>VPM3&4 x *Shi*<sup>ts1</sup> vs *Shi*<sup>ts1</sup> | 0.0005<br>0.0002 |
| 5K | One-way ANOVA<br>- Tukey post-hoc | VPM3&4 x *Shi*<sup>ts1</sup> vs VPM3&4<br>VPM3&4 x *Shi*<sup>ts1</sup> vs *Shi*<sup>ts1</sup> | 0.0088<br>0.0059 |
| 5L | One-way ANOVA<br>- Tukey post-hoc | VPM3&4 x $Shi^{ts1}$ vs VPM3&4<br>VPM3&4 x $Shi^{ts1}$ vs $Shi^{ts1}$ | 0.0254<br>0.0248 |
| 5M | One-way ANOVA<br>- Tukey post-hoc | VPM3&4 x $Shi^{ts1}$ vs VPM3&4<br>VPM3&4 x $Shi^{ts1}$ vs $Shi^{ts1}$ | 0.0045<br>0.0368 |
| **Figure S6** |  |  |  |
| S6B | Mixed-effects analysis with Geisser-Greenhouse correction and Dunnett's multiple comparison test | Odor1 vs. Odor2<br>Odor1 vs. Odor1.2<br>Odor1 vs. Odor2.2<br>Odor1 vs. Odor1.3<br>Odor1 vs. Odor2.3<br>Odor1 vs. Odor1.4<br>Odor1 vs. Odor2.4 | 0.3027<br>0.0535<br>0.0231<br>0.0145<br>0.0060<br>0.1769<br>0.3121 |
| S6C | Mixed-effects analysis with Geisser-Greenhouse correction and Dunnett's multiple comparison test | Odor1 vs. Odor2<br>Odor1 vs. Odor1.2<br>Odor1 vs. Odor2.2<br>Odor1 vs. Odor1.3<br>Odor1 vs. Odor2.3<br>Odor1 vs. Odor1.4<br>Odor1 vs. Odor2.4 | 0.1455<br>0.1078<br>0.0221<br>0.0098<br>0.0049<br>0.4316<br>0.5142 |

---

**Figure S1.**
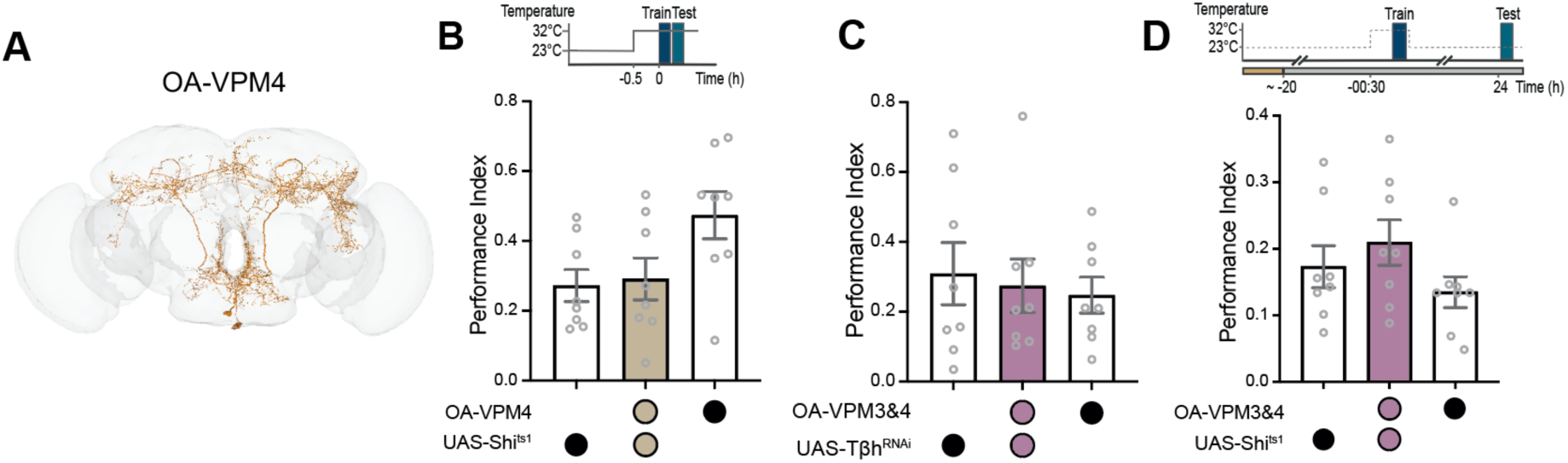
Blocking OA-VPM3 or OA-VPM4 does not disrupt STM or LTM after one training session. (**A**) FlyWire EM skeletons of OA-VPM4 neurons. (**B**) Top, temperature shifting protocol. Appetitive STM is not affected by blocking OA-VPM4 (MB113C) output during training and testing. (**C**) Top, temperature shifting protocol. Appetitive STM is not affected by expressing *Tβh*-RNAi in OA-VPM3&4. (**D**) Appetitive LTM is not affected when OA-VPM3&4 neurons are blocked with UAS-*Shi*^ts1^ during training. Data are mean ± S.E.M. Individual data points are independent biological replicates (*n*=8). Groups were compared using ANOVA with Tukey’s test for multiple comparisons.

**Figure. S2.**
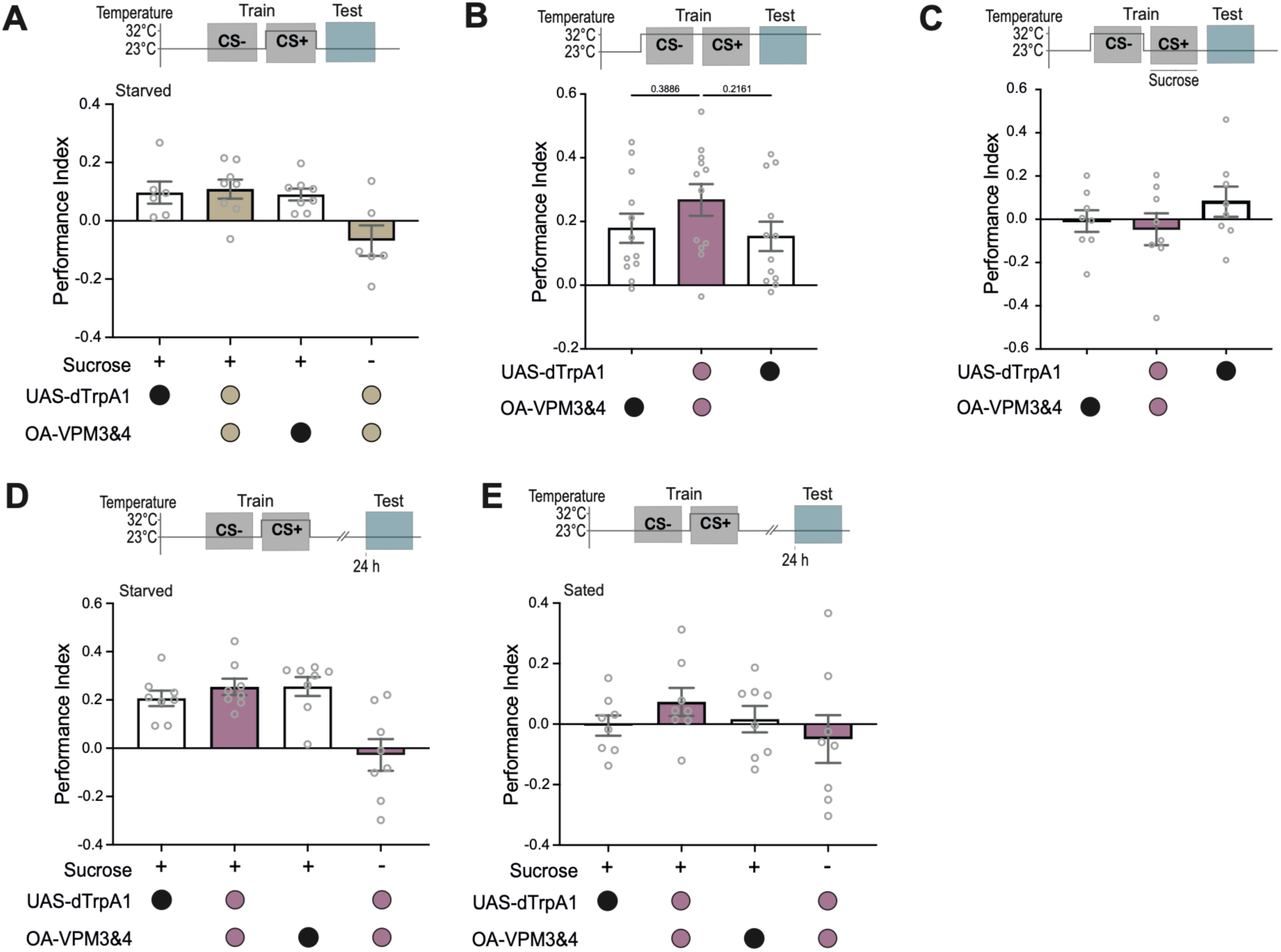
OA-VPM3 effects are specific to STM activation. (**A**-**E**) Top, temperature shifting protocols. (**A**) Thermogenetic activation of only OA-VPM4 neurons during CS+ presentation with or without sucrose presentation does not enhance appetitive STM. (**B**) STM is not statistically enhanced when OA-VPM3&4 are activated throughout appetitive training and testing. (**C**) STM is not enhanced when OA-VPM3&4 are activated with CS-exposure before CS+ with sucrose. (**D**&**E**) OA-VPM3&4 activation during CS+/sugar pairing does not enhance LTM in (**D**) starved or (**E**) sated flies. Data are mean ± S.E.M. (8≤*n*≤12). Groups compared using one-way ANOVA with Tukey’s test for multiple comparisons.

**Figure. S3.**
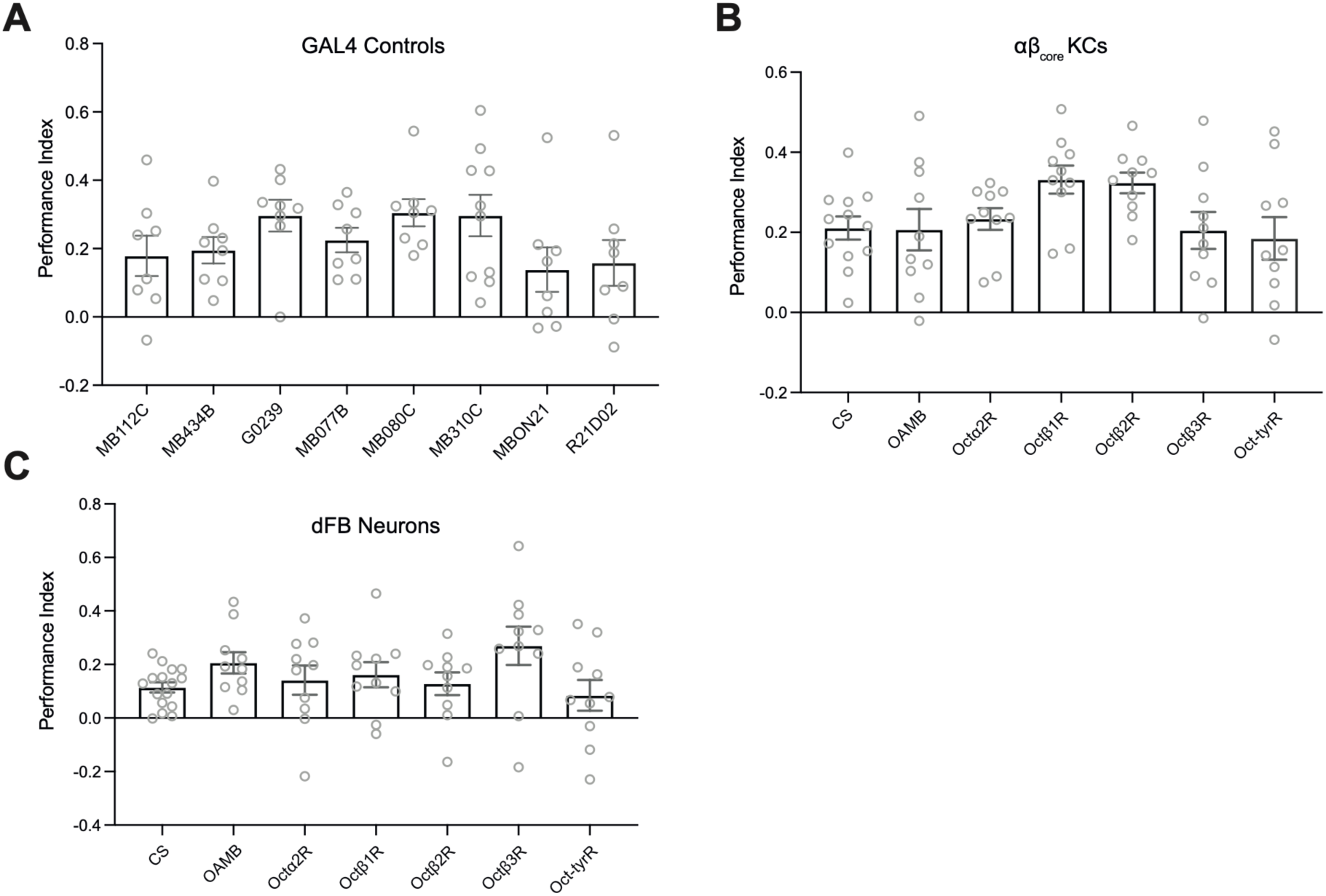
OAR knockdown in αβ_core_ KCs or dFB neurons does not affect appetitive STM. (**A**) Heterozygous MBON-GAL4 controls for the receptor RNAi screen shown in Figure. 2D-J. Appetitive STM of flies expressing OAR-RNAi transgenes in (**B**) αβ_core_ Kenyon Cells, and (**C**) a subset of dFB neurons. Data are mean ± S.E.M. (8≤*n*≤12). Groups were compared using one-way ANOVA with Dunnett’s test for multiple comparisons.

**Figure. S4.**
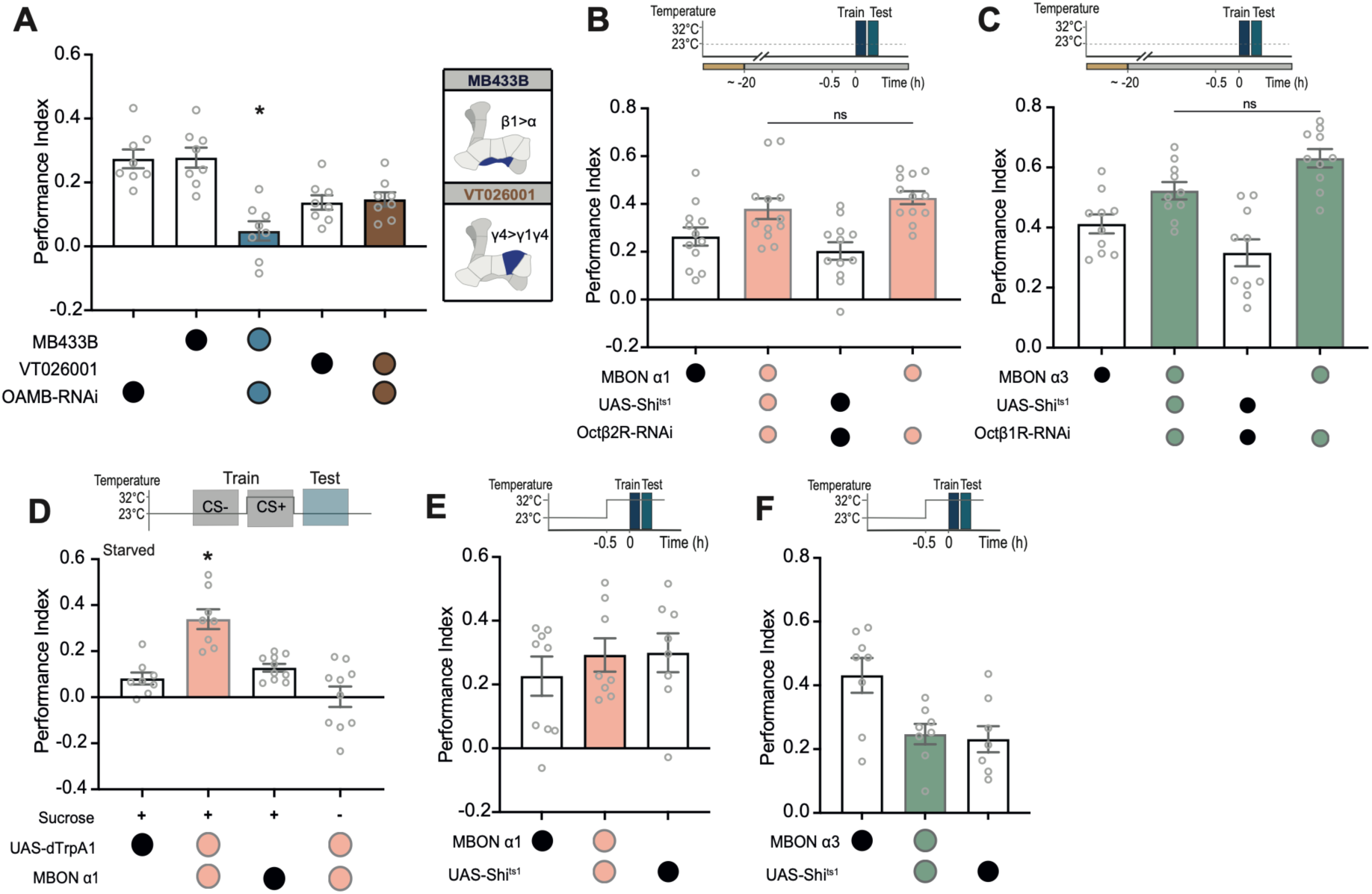
OAR-dependent modulation of β1>α, α1 and α3 MBONs is critical for appetitive STM. (**A**) Expressing *Oamb*-RNAi with MB433B (MBON β1>α), but not with VT026001 (MBON γ4>γ1γ2), impairs STM. Right, Schematic of MBONs labeled by the respective GAL4-drivers. (**B**-**C**) Top, protocol for permissive temperature control for MBON α1 and MBON α3 UAS-*Shi*^ts1^ experiments in Figure. 3G&H. STM is enhanced when (**B**) Octβ2R-RNAi is expressed in MBON α1, or (**C**) Octβ1R-RNAi is expressed in MBON α3. (**D**) Top, temperature shifting protocol. Thermogenetic activation of MBON α1 during CS+/odor pairing enhances STM, but it does not substitute for sucrose reward. (**E**-**F**) Blocking output (**E**) MBON α1 or (**F**) MBON α3 of during training and testing does not alter appetitive STM. Data are mean ± S.E.M. (8≤n≤12). Groups compared using one-way ANOVA with Tukey’s post-hoc test for multiple comparison.

**Figure. S5.**
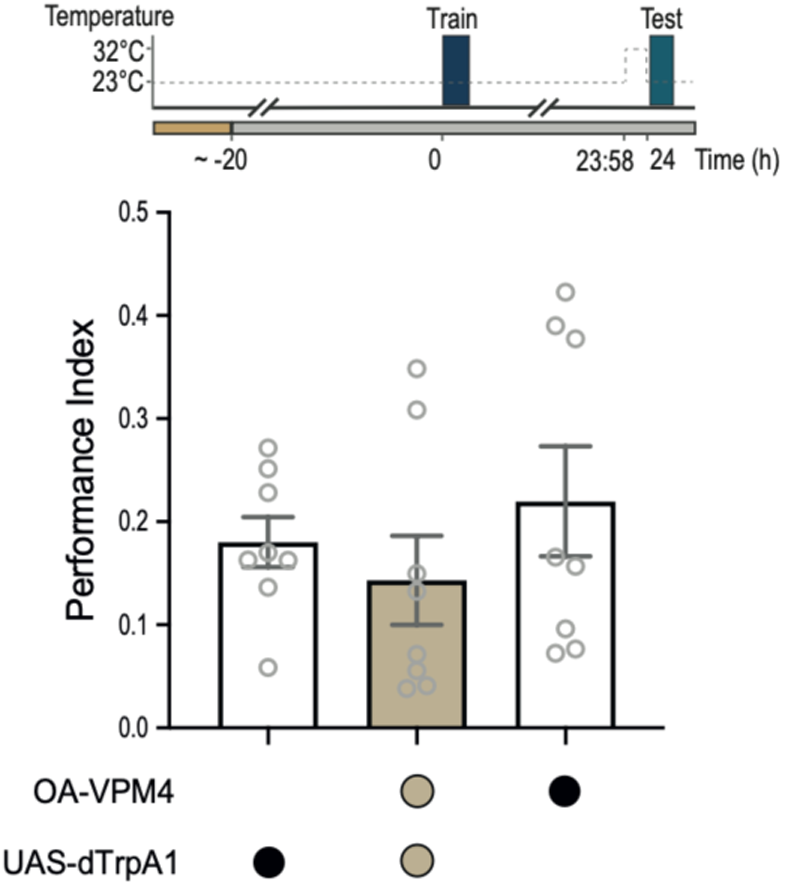
Stimulating OA-VPM4 does not impair LTM. Top, temperature shifting protocol. Activating OA-VPM4 neurons for 2 min before testing appetitive LTM has no effect on memory expression. Data presented as mean ± S.E.M. (n = 8). Groups were compared using one-way ANOVA with Tukey’s post-hoc test for multiple comparison.

**Figure. S6.**
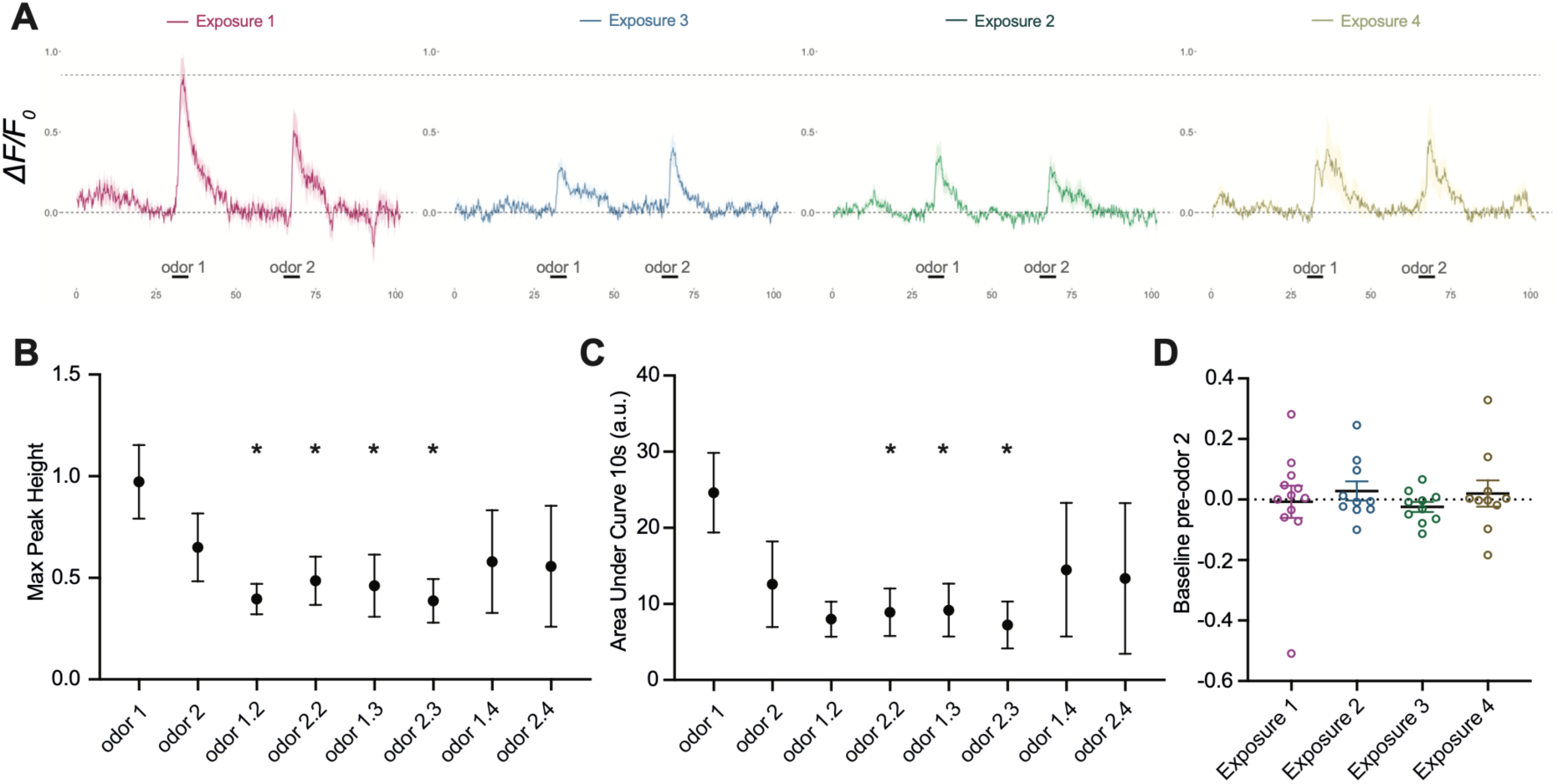
OA-VPM3 odor responses exhibit physiological adaptation. (**A**) Mean ΔF/F_0_ signal across four paired odor exposure trials. Upper dashed line indicates max peak height of the first odor response. (**B**) Mean ± SEM max peak height across all odor exposures in successive trials. (**C**) Mean ± SEM area under the curve for 10 s after odor onset for each odor exposure. (**D**) Calcium baseline signal prior to second odor exposure per trial, normalized against the baseline prior to the first exposure in that trial, indicates no change in baseline calcium signal in OA-VPM3 axons after odor exposure.

**Figure. S7.**
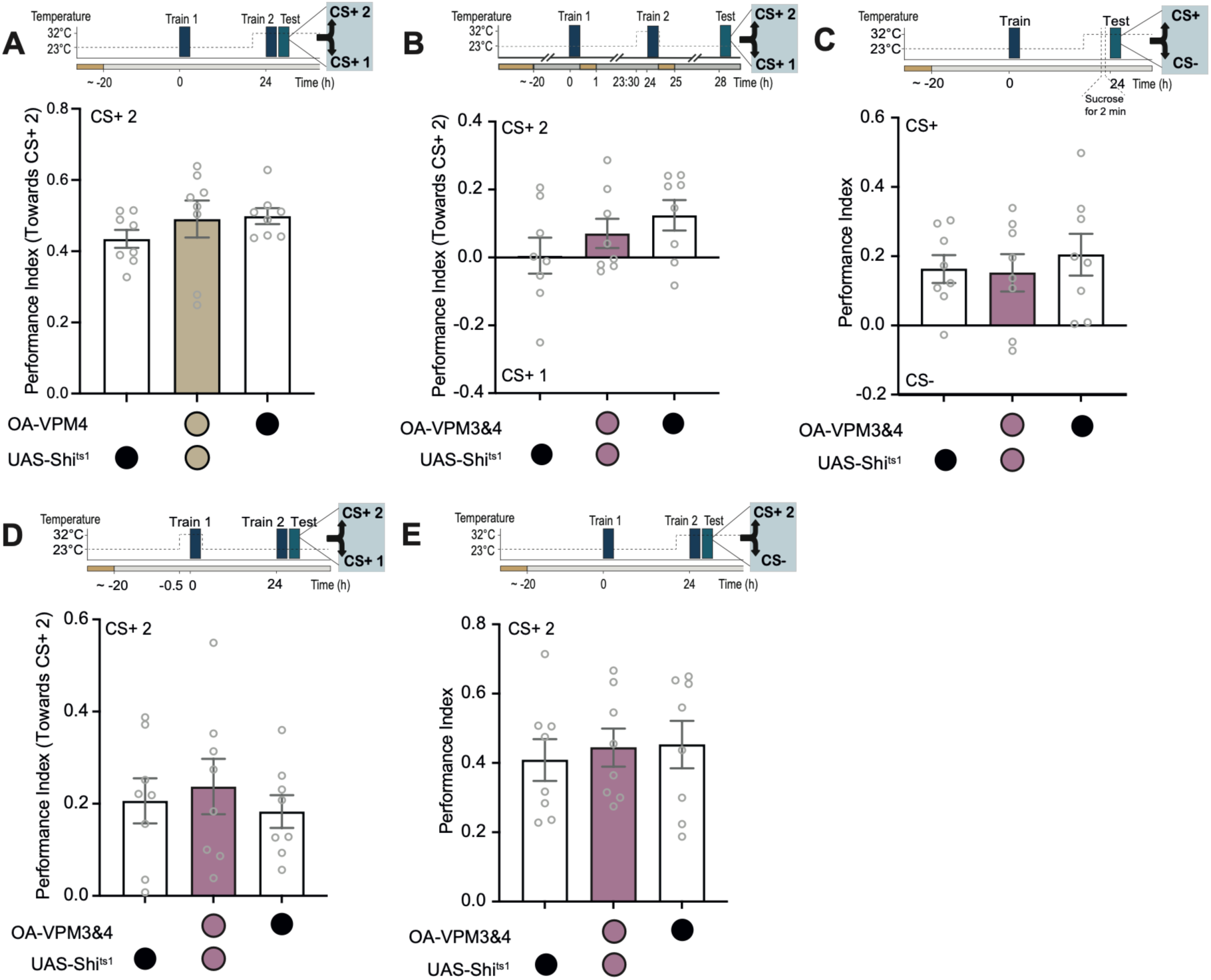
OA-VPM3 temporarily masks LTM while promoting STM. (**A**-**E**) Top, train, test and temperature shift protocols. (**A**) Blocking OA-VPM4 output during the second training and testing period does not affect preference for recent STM over remote LTM. (**B**) Blocking OA-VPM3&4 neurons during the second training does not favor expression of the oldest memory if flies are tested 24h later for 24 h CS+1 LTM vs 48 h CS+2 LTM. (**C**) Presenting only sugar for 2 min, rather than a second training session, does not make remote LTM susceptible to the enhancing effects of OA-VPM3&4 inhibition. (**D**) Blocking OA-VPM3&4 during LTM (CS+ 1) training does not affect the preference for recent CS+2 STM over remote CS+1 LTM. (**E**) Blocking OA-VPM3&4 output during recent CS+2 training and testing does not affect the preference for CS+2 over CS-odor. Data are mean ± S.E.M. (n = 8). Groups compared using one-way ANOVA with Tukey’s post-hoc test for multiple comparisons.

**Table S1.**
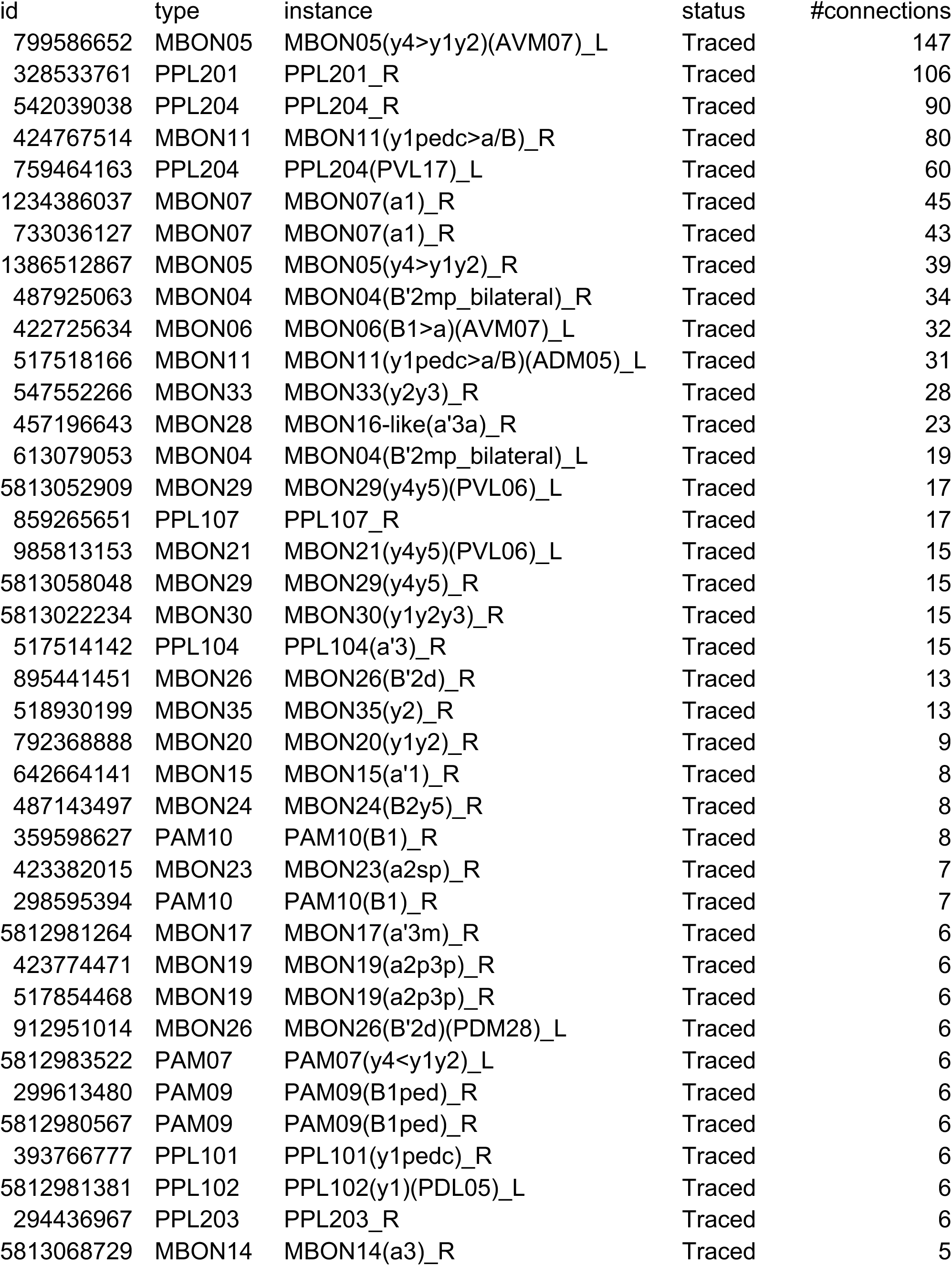
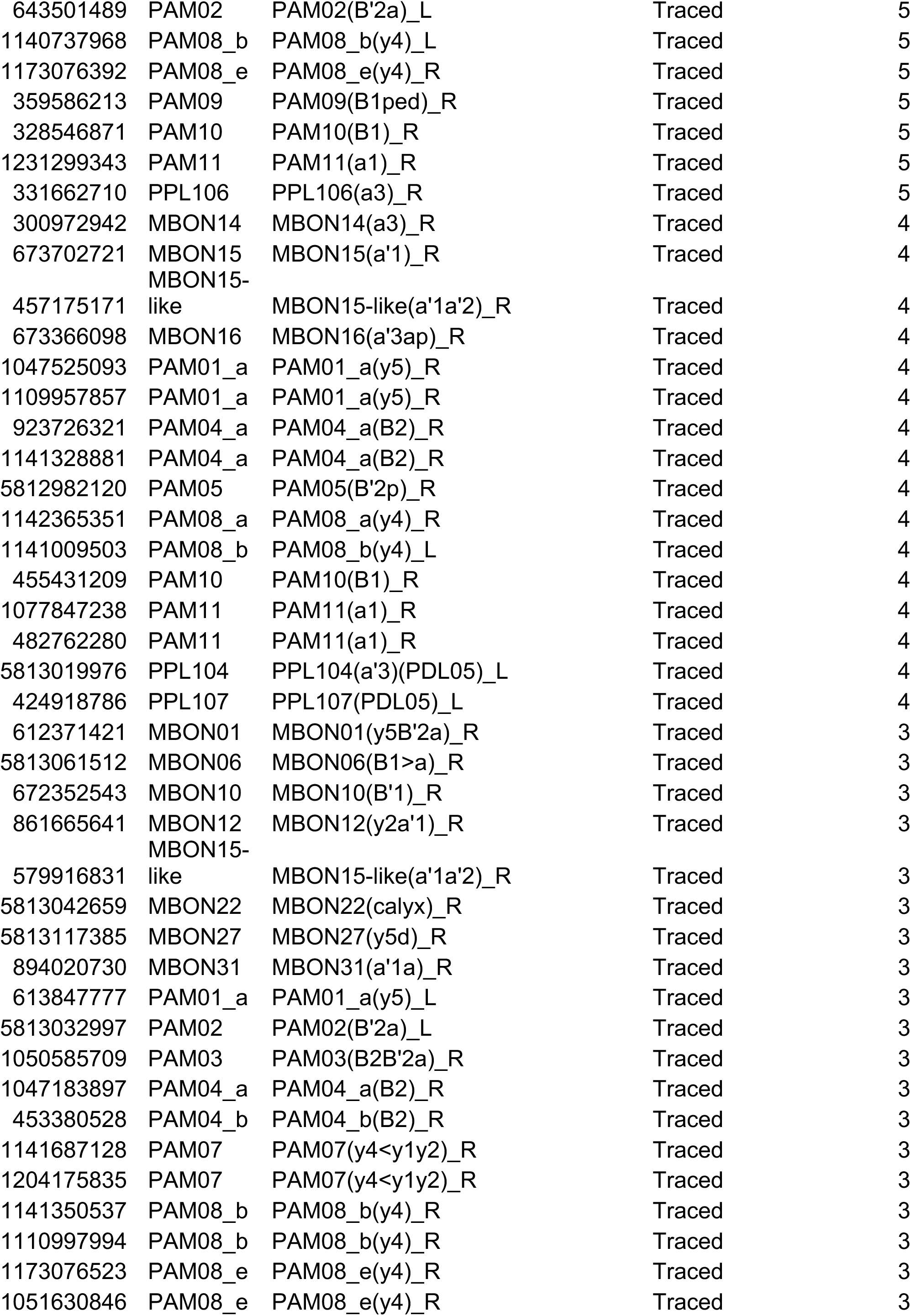
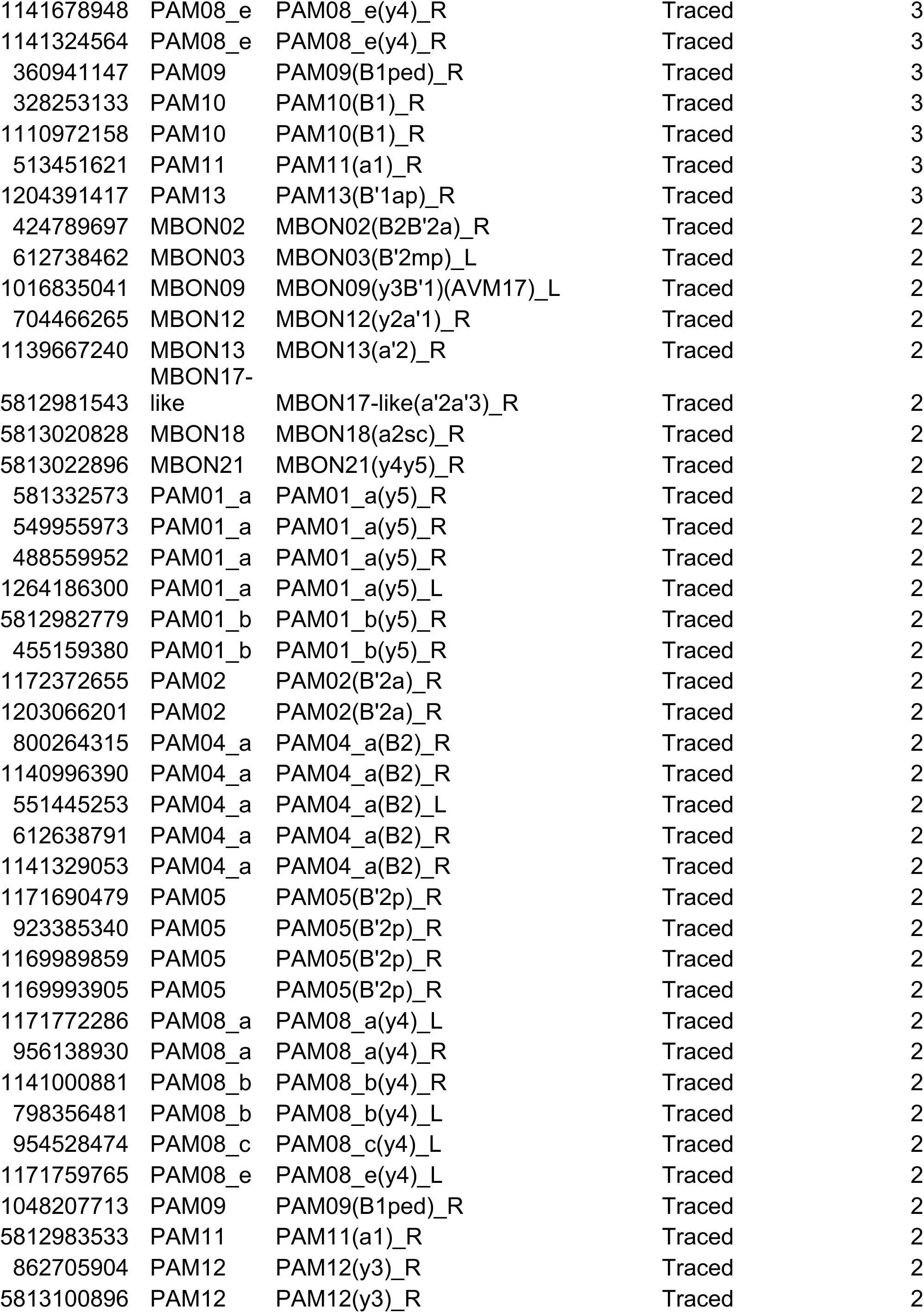
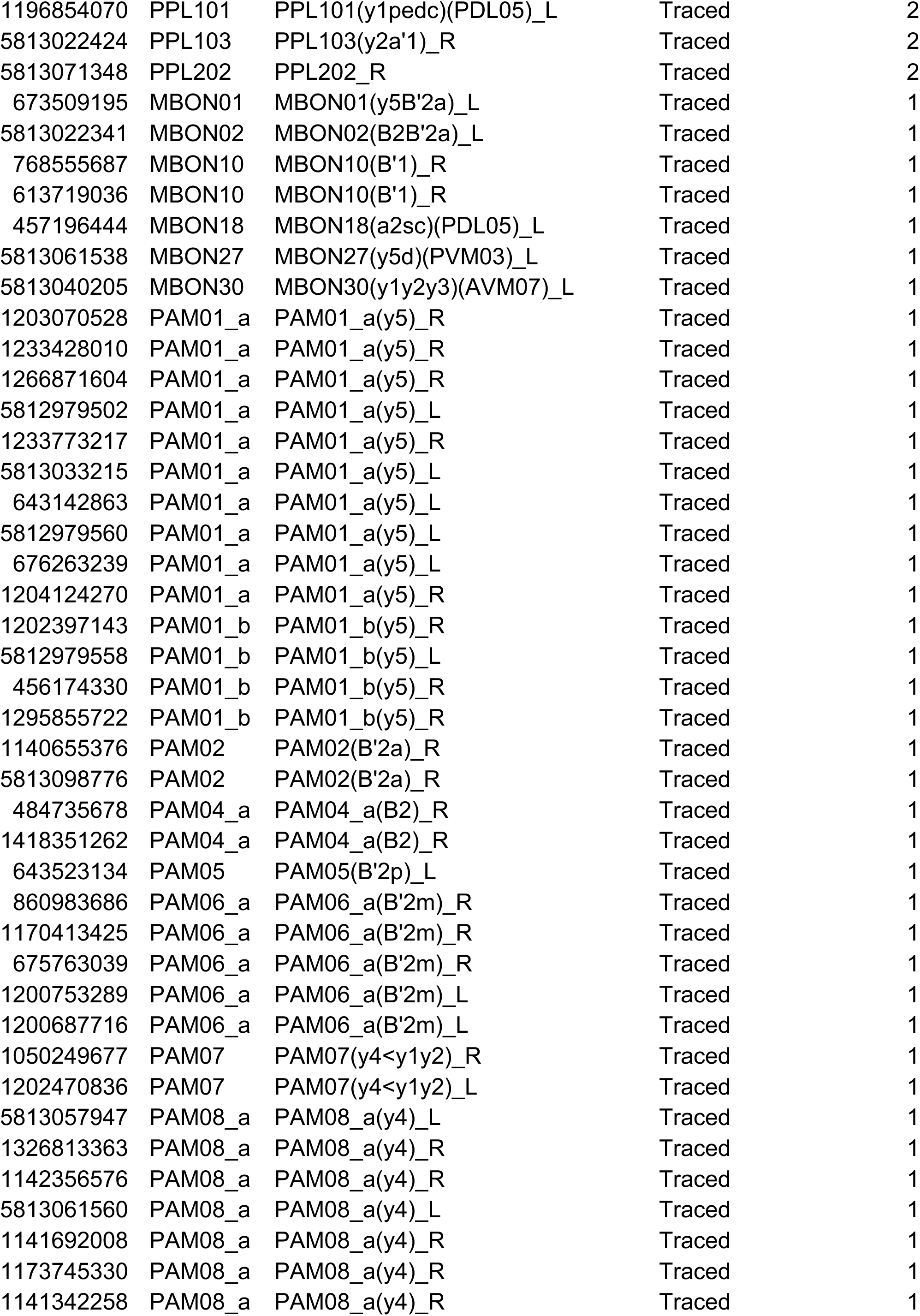
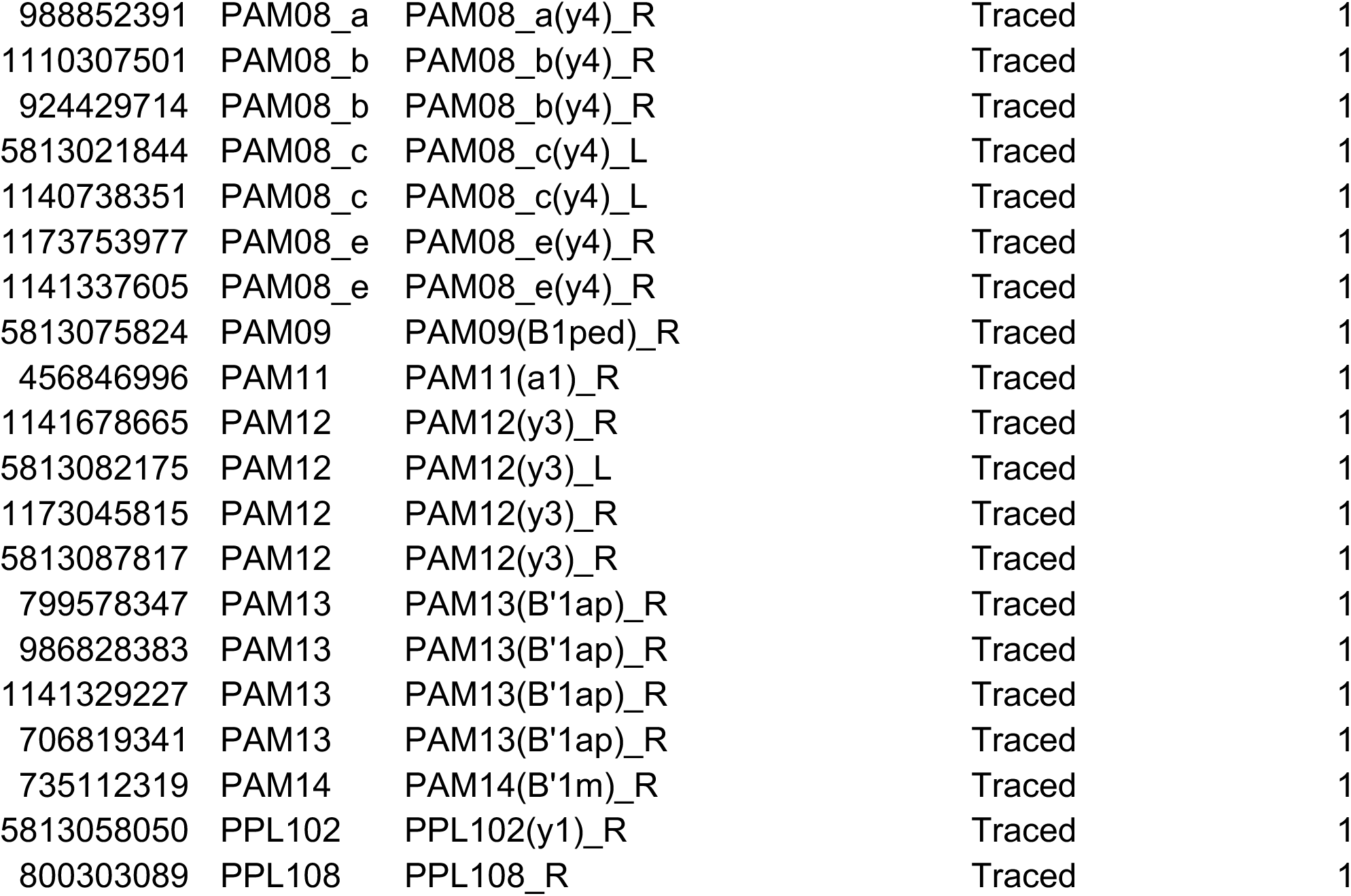
Output connectivity of OA-VPM3 (id: 329566174) to MB relevant neurons from neuprint+ (hemibrain v1.2.1)

